# Dose dependence and neurovascular mechanisms of the fMRI response to pulsed photobiomodulation in humans

**DOI:** 10.1101/2025.11.20.689588

**Authors:** Hannah Van Lankveld, Joanna X. Chen, Xiaole Z. Zhong, J. Jean Chen

## Abstract

**Background:** Transcranial photobiomodulation (tPBM) utilizes near-infrared light to penetrate the skull to stimulate neural tissue. However, the in vivo physiological response and the factors influencing this response in the human brain have yet to be understood.

**Methods:** In this study, we utilize functional magnetic resonance imaging (fMRI) to evaluate the effect of tPBM on the blood-oxygenation (BOLD) and cerebral blood flow (CBF), while varying stimulation parameters such as wavelength, irradiance, and frequency. We further examine the influence of skin tone and sex. We further model the neurovascular interactions underlying the response.

**Results:** Our results show that the fMRI responses to tPBM is not restrained to the site of irradiation, but quickly spreads to distal sites. Certain regions display an fMRI response sustained after tPBM cessation. Importantly, the responses are dependent on biological and stimulation parameters. Lastly, biophysical modeling revealed a consistent neurovascular coupling-like behaviour underlying these responses.

**Conclusion:** Empirical characterizations of dose dependence are critically important to brain stimulation methods in general but have yet to be demonstrated in most cases. This is the first tPBM study to do just that, establishing the foundation for precision medicine using tPBM, and sets a valuable precedent for the field of brain stimulation.

## Introduction

Transcranial photobiomodulation (tPBM) is a non-invasive neuromodulation technique that utilizes low-level red or near-infrared (NIR) light [1]. NIR light can penetrate human skin and tissue to various depths ranging several millimeters, giving rise to biological reactions [2][3]. Specifically, tPBM is known for its ability to increase adenosine triphosphate (ATP) production through the dissociation of an inhibitory nitric oxide (NO) molecule on the cytochrome c oxidase (CCO) enzyme [4], [5]. tPBM is currently in clinical trials for the management of stroke, Parkinson’s disease, depression, Alzheimer’s disease and other dementias [6]. Recent studies have shown the various positive cognitive effects of tPBM, such as enhancing memory processing [7], improving executive function [8], as well as enhancing cognitive and emotional brain functioning [9]. While these effects have been well distinguished in-vitro, and through cognitive testing, it is important to recognize that individual cellular responses can propagate into broader physiological changes in vivo. Upon photon absorption, the NO molecule is photodissociated from CCO, as NO is a potent vasodilator, its release can directly influence vascular tone, cerebral blood flow (CBF) and potentially neurovascular coupling. These secondary vascular responses cannot be fully inferred from cellular models or cognitive testing alone; rather they require direct in vivo measurement to understand how PBM shapes cerebrovascular physiology and brain function. Now that we know tPBM modulates CSF flow [10], an important question is how this occurs, and whether such effects are mediated by vascular or metabolic pathways. Despite the growing evidence supporting tPBM as a neuromodulation therapy, the physiological mechanisms of tPBM on the human brain remain poorly understood, as tPBM could influence metabolism, oxygen consumption, and perfusion [11].

To assess the physiological effects of tPBM on the human brain, several neuroimaging techniques have been employed, including single-photon emission computed tomography (SPECT) [12], functional near-infrared spectroscopy (fNIRS), electroencephalography (EEG), and blood-oxygenation-level-dependent (BOLD) functional magnetic resonance imaging (fMRI). Among these, fNIRS has been the most frequently used in real-time PBM studies, showing time-locked hemodynamic changes at the site of stimulation [13]. fNRIS is portable and has a quick setup time, but its limited spatial coverage restricts whole-brain interpretation [14]. fMRI offers whole-brain coverage and sensitivity to both hemodynamic and metabolic changes [15], making it well-suited for quantifying tPBM-induced effects on cerebral physiology. Nonetheless, to our knowledge the majority of fMRI studies in PBM have examined the pre-post-PBM differences [16] [17] [18], and little is known about the real-time cerebrovascular effects, as prior simultaneous PBM-fMRI studies have not assessed direct CBF or BOLD time series responses [19], [20].

BOLD fMRI measures changes in deoxygenated hemoglobin concentrations, which reflect the balance between the cerebral metabolic rate of oxygen (CMRO₂) and CBF. Neural stimulation increases CMRO_₂_, accompanied by vasodilation and a corresponding rise in CBF [21]. However, the interpretation of BOLD with regard to CBF and CMRO_₂_ is ambiguous, and thus combining BOLD and CBF measurements provides complementary insights. tPBM can increase CBF through nitric oxide induced vasodilation, distinguishing the rise in blood flow from oxygen metabolism, and the overarching BOLD response provides the full picture of the hemodynamic response to tPBM. The literature on this topic has been varied. While fNIRS studies of the frontal cortex have widely shown an increase in the concentrations of oxyhemoglobin and total hemoglobin (a surrogate of CBF) due to tPBM [14], a 2021 fMRI study showed that an 810nm, 250mW/cm^²^ irradiation pulsing at 10Hz did not induce a significant CBF change [18]. A more recent study showed a significant increase in CBF and CMRO_₂_, but the analysis was limited to the site of irradiation [22]. In fact, beyond the question of the BOLD and CBF responses in tPBM, there remain many significant gaps in the understanding of how tPBM affects neural activity and vascular responses. For example, is the BOLD response to tPBM focal to the site of irradiation or more global? How is the BOLD response during tPBM shaped by changes in CBF? Addressing these questions is crucial to broadening the understanding of the effect of PBM on the brain.

Beyond establishing the neurovascular underpinnings of tPBM, it is also important to determine the dose dependence. The cerebrovascular response may depend on specific stimulation parameters such as wavelength, irradiance, and pulsation frequency. Shorter wavelengths have been shown to penetrate less deeply than longer NIR wavelengths [23], [24], [25], [26]. Wavelengths around 670nm and 830nm are known to correspond to absorption peaks of CCO, the targeted photo-acceptor, but wavelengths in the range of 808nm to 1064nm are typically used. Irradiance, the energy of light delivered per unit area (mW/cm^²^), directly influences energy accumulation in targeted brain regions. Typical irradiance settings for clinical applications range from 250 to 300 mW/cm^²^, staying below the medical safety limit of 330 mW/cm^²^. Pulsation frequency, specific to pulsed tPBM, is typically set to 10Hz or 40Hz to align with the brain’s natural alpha and gamma rhythms, respectively, but their effects have yet to be distinguished in vivo. Furthermore, biological factors such as melanin, which also impacts penetration [27], has not been considered in tPBM.

In this study, we aim to assess data-driven region-specific and real-time BOLD and CBF responses to varied doses of pulsed tPBM in healthy adults. Leveraging simultaneous tPBM-fMRI measurements of BOLD and ASL derived CBF, we quantify how different stimulation doses shape the temporal profile and enable us to identify regions of peak sensitivity, examining both vascular and metabolic components. To our knowledge, this is the first study to systematically investigate the dose-dependent effects of wavelength, irradiance, frequency, skin tone and sex on real-time BOLD and CBF responses, making this the first systematic, multi-parameter in vivo neurovascular investigation. Moreover, based on prior literature and our own simulation studies, we hypothesize that individuals with lighter skin will experience a greater percentage change in BOLD and CBF than those with intermediate or darker skin tones. Additionally, we hypothesize that tPBM will induce a larger percentage change in BOLD and CBF with the 808nm wavelength compared to 1064nm. We will also determine the dependence of the tPBM response and the underlying neurovascular coupling on light-pulsation frequency as well as irradiance and sex.

## Methods

### Participants

We recruited 45 healthy young adults (age 20-32, 23 M/22 F) who were screened prior to their participation to ensure there was no history of neurological or physiological disorders, malignant disease, or the use of medications that could have influenced the study. The Baycrest Research Ethics Board (REB) approved the study, and all experiments were conducted in accordance with REB guidelines, with each participant providing written informed consent. The sample size was calculated pre-acquisition, using the P, W, CI calculation. A 95% confidence interval, an expected proportion of p=0.8 and a total width of 2× standard deviation (SD=0.3) was inputted. Based on these expected values, the output sample size was 32, when using the observed 2x standard deviation of 0.4, the corresponding sample size was 17.

### PBM Instrumentation

The MRI-compatible lasers were provided by Vielight Inc. (Toronto, Canada), and the light was delivered through a 10-meter, 400-μm diameter optic cable to the MRI, while the laser system remained outside the scanner room. The light parameters, including wavelength, irradiance and pulsation frequency, were controlled from the scanner console while the participant was inside the scanner. To avoid any placebo effects, the participant remained blinded to the stimulation paradigm, the light was secured through a headpiece such that it was not visible to the participants.

### MRI Acquisition

MRI data were acquired using a Siemens Prisma 3 Tesla System (Siemens, Erlangen, Germany) with a 20-channel radio-frequency head coil. For each participant, the following data was collected: (a) A T1-weighted structural image (sagittal, 234 slices, 0.7 mm isotropic resolution, TE = 2.9ms, TR = 2240ms, TI = 1130ms, flip angle = 10°). (b) A dual-echo pseudo-continuous arterial spin labeling (DE-pCASL) scan (courtesy of Danny J. J. Wang, University of Southern California) to simultaneously measure CBF and BOLD (TR = 4.5s, TE1 = 9.8ms, TE2 = 30ms, post-labeling delay = 1.5s, labeling duration = 1.5s, flip angle = 90°, 3.5mm isotropic resolution, 35 slices, 25% slice gap, total scan time = 12 minutes 15 seconds). Additionally, an M0 calibration scan was acquired for CBF quantification, with a TR of 10 s and a scan time of 40 s, while all other parameters remained the same as in the pCASL scan. For n=30 participants, one MR thermometry sequence was acquired (TR = 97ms, TE1 = 4.92ms; TE2 = 7.38ms; FoV = 192mm; slice count = 10).

Participants were scanned while viewing naturalistic stimuli to minimize variations in brain state across participants, thereby improving the reproducibility of the stimulation outcome (Gal et al., 2022).

### Stimulation Protocol

The laser parameters varied across two wavelengths, three irradiances and two pulsation frequencies, as shown in **Table 1**. Moreover, we recruited equal representations of three skin colour groupings (light, intermediate and dark). As a quantitative indicator of skin pigmentation (eumelanin and pheomelanin), the individual typology angle (ITA) of each participant was measured using a CM-600D Spectrophotometer (Konica Minolta). The calculated ITA and pixelated image of the right forehead for each participant is shown in **Figure 1**.

**Figure 1.**
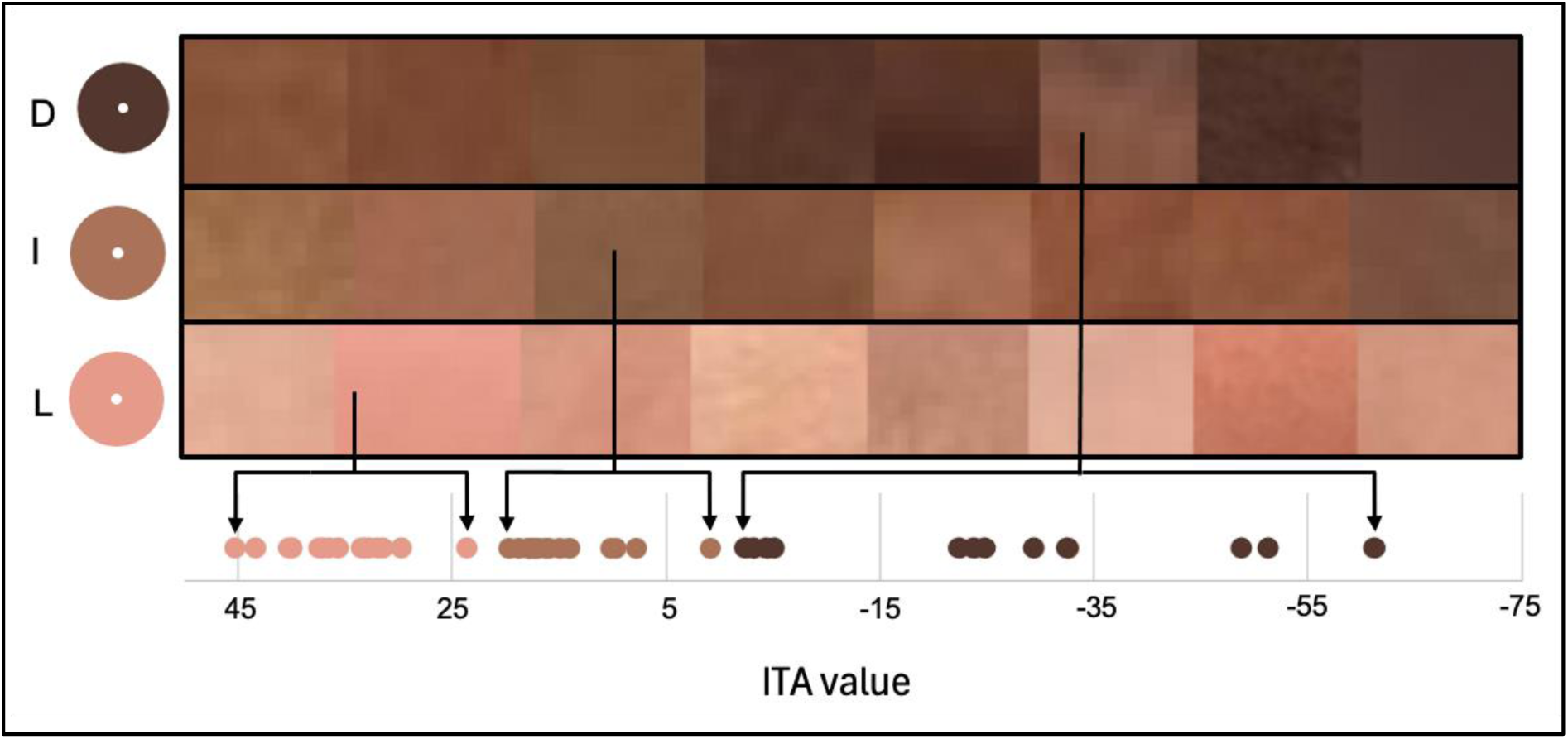
Objective measurements of skin tone as compared to photographs of forehead skin for a representative group of subjects. The images were categorized based on their individual typology angle (ITA), to show the general tone of each skin tone grouping, i.e. light (L), intermediate (I) and dark (D). The dots represent the actual ITA values corresponding to the mean visually perceived skin tone based on the individual photos of skin patches which are ordered by ITA in each skin-tone grouping. That is, there are differences between visually quantifying skin tone and the representation through ITA. A higher ITA reflected lighter skin. Additional details can be found in the Supplementary Materials section **S.1.2**.

**Table 1:**
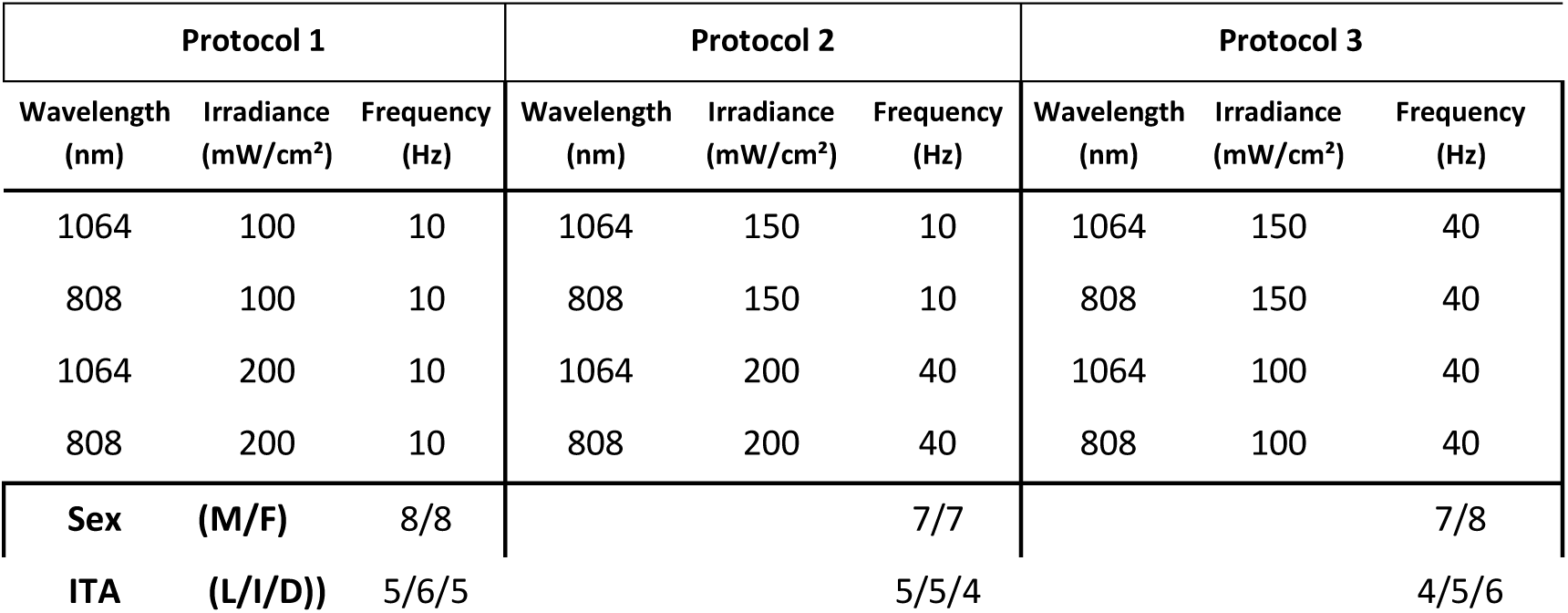
Stimulation paradigm for each protocol grouping. . The male (M) to female (F) and skin-tone distributions are similar across the three groups. L: light; I: intermediate; D: dark skin. Each subject underwent 4 tPBM sessions each with a unique combination of dose parameters.

In addition to skin tone, another biological factor to consider is the distance from the skull to the cortical surface, which can differ between sexes and individuals. To quantify this measurement, the T1-weighted structural MRI scans for each participant were acquired. The Freesurfer measurement tools were used to estimate the distance from the skull to the cortical surface.

The tPBM laser was carefully positioned on the right side of the forehead (**Figure 2**), targeting the right prefrontal cortex, with two vitamin E capsules placed as fiducial markers of the precise location of irradiation. Each participant was assigned to one of three protocols, with each protocol comprising four scans (see **Table 1**), resulting in 15 participants under each protocol.

**Figure 2.**
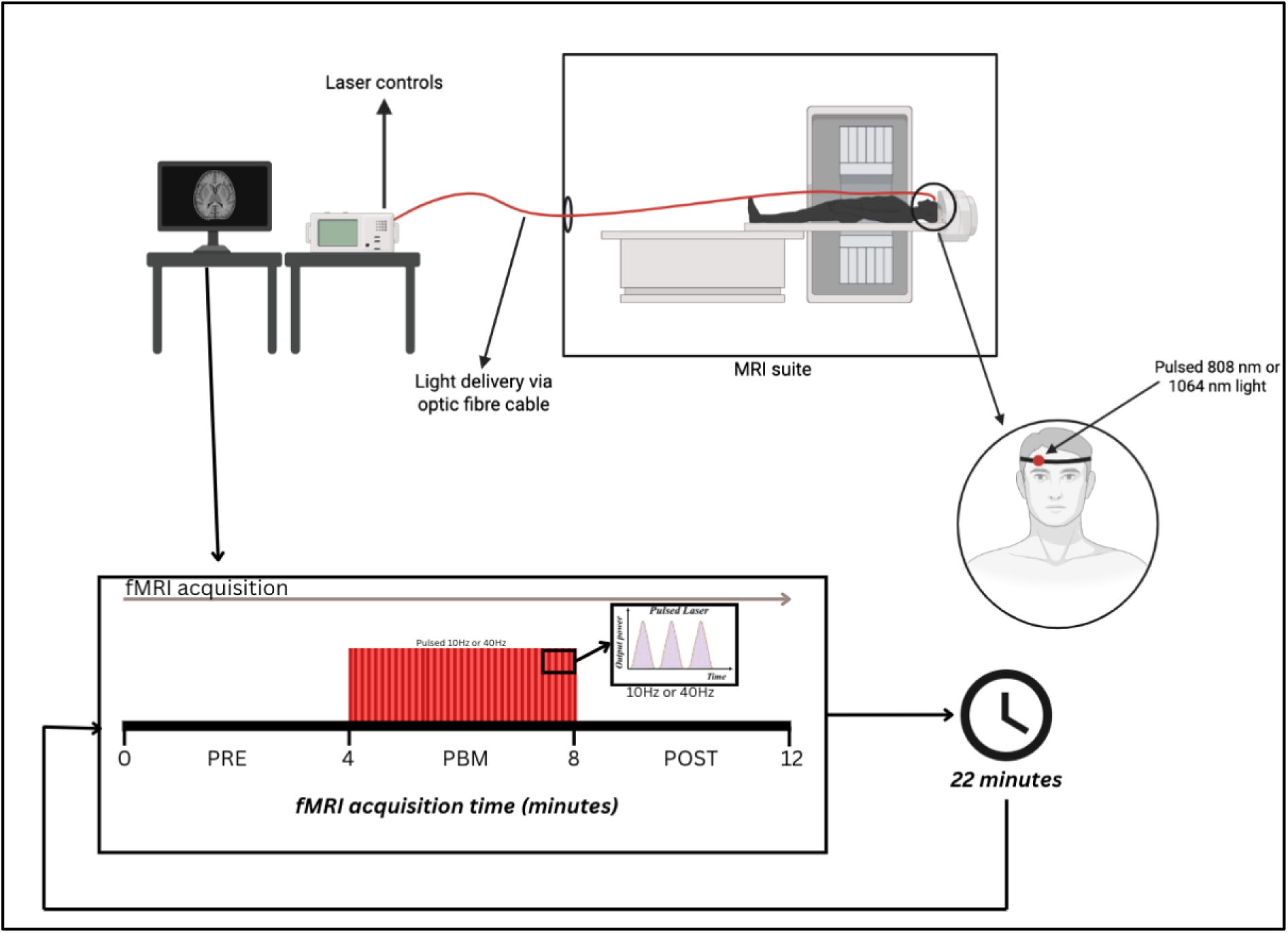
tPBM-fMRI experimental setup. The optic fibre cable carries the NIR light from the laser control system through a small hole into the MRI suite. The optic fibre is secured to the participants chest strap, fed through the RF coil in the MRI and positioned on the right side of the forehead. From outside the MRI suite, the controls at the console allow the laser to be turned ON/OFF and delivered at the prescribed parameters throughout the fMRI acquisitions. fMRI acquisition follows a 4min-OFF, 4-min-ON, 4-min-OFF paradigm.

The stimulation paradigm was 4-min-OFF, 4-min-ON, 4-min-OFF, as shown in **Figure 2**. A break of at least 10 minutes was ensured before each subsequent tPBM session. In combination with the 4 minutes of pre-stimulus acquisitions and 4 minutes of post-stimulus acquisitions, there was a minimum time of 18 minutes between stimulations.

### MR Thermometry

To test for the existence of brain-temperature increases during tPBM, MR thermometry data were collected for the sessions corresponding to the 1064 nm wavelength delivered at the highest irradiance (200 mW/cm^²^) at both 10Hz and 40Hz (N =30 participants). The full details are provided in Supplementary Materials.

### Data Preprocessing

BOLD-fMRI preprocessing was performed using a custom script that employed tools from the FMRIB Software Library (FSL) [28], AFNI [29], FreeSurfer [30] and MATLAB. The pCASL datasets were brain extracted (FSL bet) and split into control and tag images (FSL slicetimer and FSL fslsplit), motion correction was done separately for each. The BOLD data were generated using the surround-averaging of control and tag frames [31]. CBF-fMRI data was preprocessed using ASLprep [32]. MATLAB was used for temporal and spatial outlier removal, drift correction and normalization. The first five volumes of both BOLD and CBF were rejected to allow the fMRI signal to enter a steady state. All datasets were registered and resampled into MNI space for group analysis.

### fMRI analysis

Following preprocessing, the data was input into FSL MELODIC [33] for an model-free independent component analysis (ICA) regardless of protocol and melanin grouping (N = 180 data sets). The ICA was chosen over the conventional general-linear model analysis, as the hemodynamic response function for tPBM remains unknown. The number of independent components (ICs) was set to 30 to avoid fragmentation. The ICs were normalized, and their temporal characteristics were examined through a t-test to determine significant correspondence with the onset of the tPBM stimulus. Subsequently, dual regression was performed to map the ICs to the normalized individual subject time series. The dual-regression model consists of three stages, generating statistical maps and time series for each IC, allowing for subject-specific characterization of BOLD dynamics and spatial specificity. Each of these ICs was taken as a region-of-interest (ROI). After dual regression, the subject-specific BOLD-normalized and CBF-normalized time series for each ICA-generated ROI plus the site of irradiation were plotted to assess the dynamics of BOLD and CBF signals across stimulation conditions.

### Mixed-Effects Modeling

The t-scores associated with the %BOLD and %CBF changes were entered into a linear mixed-effects (LME) model as the dependent variable to analyze the influence of variables including irradiance, wavelength, frequency, melanin level (ITA), sex and cortical distance on the fMRI responses. However, if the time series did not exhibit a statistically significant change, the %BOLD and %CBF change for that condition was set to zero. The LME model followed a two-step approach. First, we used MATLAB’s stepwise LME (stepwiselm) function to rule out variables that had no significant effect (p > 0.05, False Discovery Rate (FDR)-corrected) on the temporal BOLD and CBF response. These variables were excluded from the final LME to maximize information content of the remaining fixed effects. The final model followed the following format, where the subject ID was included as a random variable.

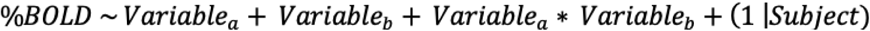

Wavelength, irradiance, pulsation frequency and sex were set as categorical variables, and ITA as a continuous variable. Final statistical p-values were thresholded with FDR correction.

### BOLD Biophysical Modeling

To further characterize the relationship between BOLD and CBF responses specific to tPBM, we fit the temporal-mean %BOLD and %CBF responses from each scan of each subject to the deoxyhemoglobin dilution model, as shown in Eq. 1 [34], with set parameters β=1.3, α=0.2 [35,36]. The relationship between CBF and CMRO₂ was defined by Eq. 2 [34,37],

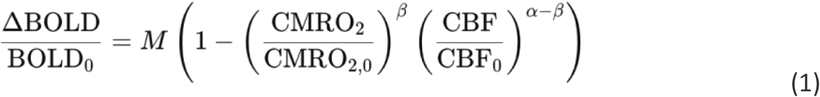

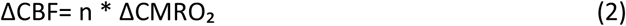

where changes in CBF and CMRO₂ are related by the neurovascular coupling ratio n. The fits for n and M were performed on different windows of the BOLD and corresponding CBF responses, for each ROI separately, where the M-value estimated from the first-time window was then used in the two subsequent windows to fit for the n-values. To reduce noise and stabilize the CBF-BOLD relationship prior to model fitting, the data was stratified into bins of 6 points.

## Results

### MR thermometry

Group-level statistical comparison against the pre-stimulus (minutes 1-4) and the stimulus (minutes 5-8) was conducted to show that the average temperature change was minimal and remained stable over the entire 12 minutes **(Table S2).** For 10 Hz stimulation, the mean temperature increased from - 0.02 +/- 0.18°C during PRE to 0.05 +/- 0.17 °C during PBM, but this change was not statistically significant (p = 0.13). Likewise, for 40 Hz stimulation, the temperature increased from -0.01 +/- 0.16 °C to 0.01 +/- 0.15 °C, with no significant difference observed (p=0.76). The results confirmed that even at the longest wavelength and highest irradiance used (corresponding to the highest energy deposition used), no measurable thermal effects were produced in the illuminated brain region **(Table S2)**. Please see **Figure S2** in Supplementary Materials for details on the definition of the illuminated region.

### BOLD fMRI responses

The ICA revealed 3 ROIs that were associated with the tPBM timing, based on visual inspection. Additional information regarding the ICA and dual regression approach can be found in the Supplemental Materials section.

Three ICs survived the ICA step and are shown in **Figure 3**, with the anatomical and functional determinants of ROIs 1-3 obtained from NeuroSynth (correlation coefficients (r) listed below) [38]:

- ROI 1 (**Figure 3a**) consists of subgenual (r = 0.55), accumbens (r = 0.45), and ventromedial prefrontal (r = 0.43), which are associated with mood regulation and emotional processing;
- ROI 2 (**Figure 3b**) consists of the posterior superior (r = 0.31), the temporal (r = 0.30), and the temporal sulcus (r = 0.30), which are involved in information processing;
- ROI 3 (**Figure 3c**) consists of the posterior cingulate (r = 0.56), the precuneus (r = 0.46), and the default mode network (DMN) (r = 0.35) which are associated with cognitive functioning, emotional regulation, and memory.
- ROI 4 (**Figure 3d**) represents the illuminated region, defined as the region most proximal to the

light’s incidental site, as described earlier and in **Figure S2**.

**Figure 3.**
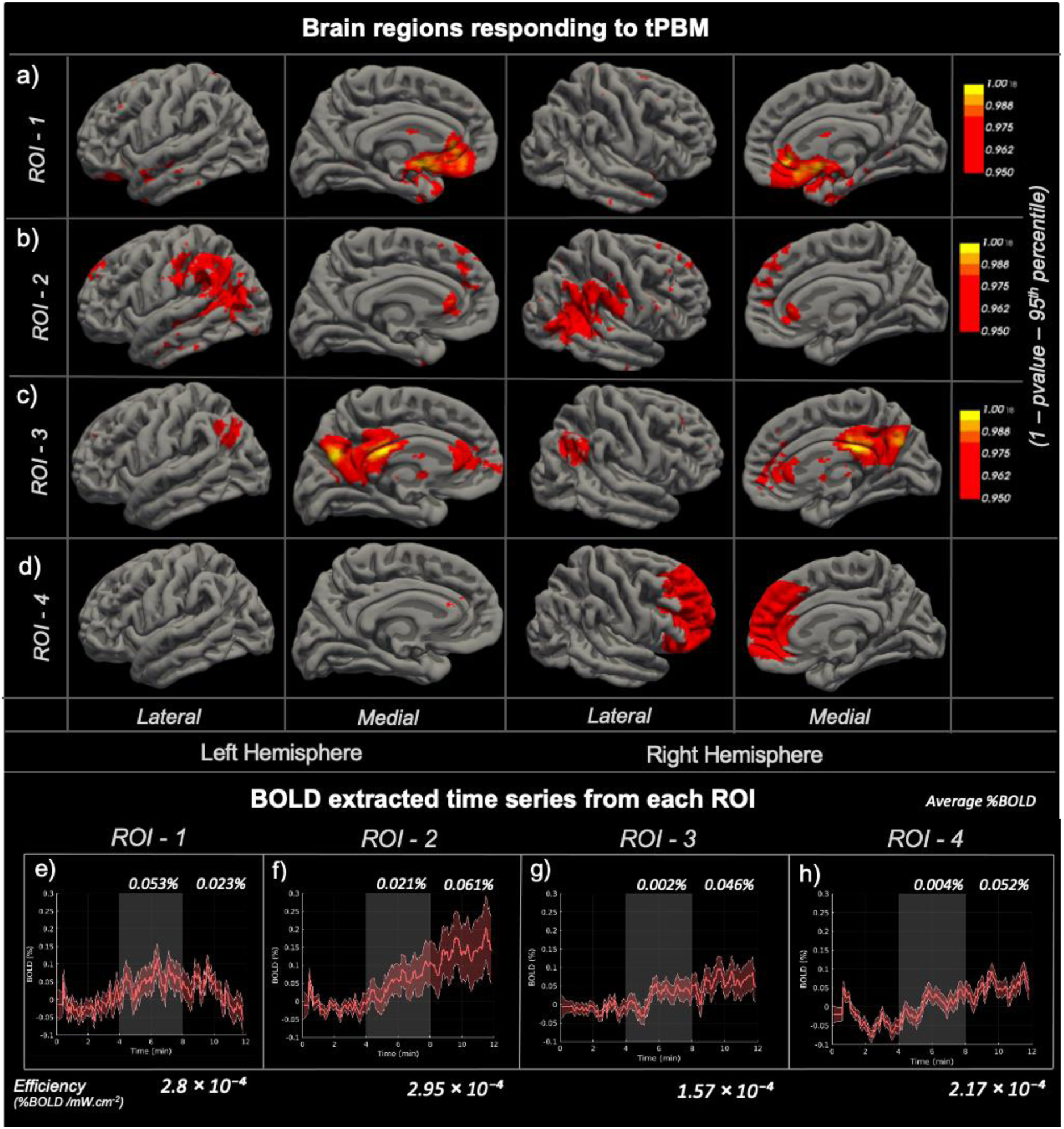
**Overview of the brain regions responding to tPBM. (**a-c) ROIs 1-3, which are Independent components showing a significant BOLD fMRI response irrespective of subject or stimulation parameter, thresholded to p < 0.05. (d) ROI 4, which is the site of stimulation (irradiation), manually selected per participant (see Supplemental Materials **S.1.3**) and averaged across all subjects. All ROIs are mapped from native to MNI space, and shown on the medial and lateral surfaces. Only voxels with t-values corresponding to p < 0.05 are displayed, the colour bar represents the thresholded t-values from the stage 3 dual regression analysis. The colourbars represent 1 minus the p-value, with a value of 1 indicating the strongest significance of the BOLD response t-score. **Mean BOLD time series from each ROI defined in a-d;** To characterize the temporal responses of each ROI, we extracted the BOLD signal trajectory in each ROI by computing the average BOLD intensity across all active voxels. Coloured shaded regions represent the standard error across all datasets. The tPBM stimulus “on” period (minutes 4:8) is denoted by the shaded grey region. The mean %BOLD increase during stimulation and post stimulation is noted for each ROI along with a metric of efficiency, determined by normalizing the largest %BOLD response by irradiance, these values are also tabulated in **Table S3**.

### Linear Mixed Effects Modelling

The LME modelling highlighted the ROIs exhibiting a significant temporal response magnitudes between various stimulation parameters, a summary of all responses are shown in **Table 2** below.

**Table 2.** ROI-level statistical summary results from the Linear Mixed Effects model of stimulation parameters for both a) BOLD and b) CBF responses. ITA breakdown: L/I/D = light, intermediate, dark). T-test describes the statistical comparison performed, either between stimulation (PBM) and pre-stimulus baseline (Pre) or between post-stimulation (Post) and the pre-stimulus baseline (Pre). Normalized Effect Size provides the magnitude of the response change after normalization to the baseline across all datasets. The detailed statistical information can be found in S.4 in Supplementary Materials.

a) %BOLD
| ROI | Location | T-test | Effect | Significance FDR corrected p value | Normalized Effect Size |
| --- | --- | --- | --- | --- | --- |
| 1 | Subgenual /<br>Accumbens | Pre-PBM | Wavelength<br>1064 > 808 (nm) | p = 0.006 | +0.15 |
| 2 | Posterior Superior /<br>Temporal Sulcus | Pre-Post | Wavelength<br>1064 < 808 (nm) | p = 0.041 | -0.07 |
| 3 | Posterior Cingulate /<br>DMN | Pre-Post | Wavelength<br>1064 < 808 (nm) | p = 0.033 | -0.12 |
| 1 | Subgenual /<br>Accumbens | Pre-PBM | Irradiance<br>100 < 150 > 200 (mW/cm<sup>2</sup>) | p = 0.045 | +0.09 |
| 3 | Posterior Cingulate /<br>DMN | Pre-PBM | Irradiance<br>100 < 150 < 200 (mW/cm<sup>2</sup>) | p = 0.041 | +0.21 |
| 4 | Stimulus Site /<br>Prefrontal Cortex | Pre-Post | Irradiance<br>100 > 150 < 200 (mW/cm<sup>2</sup>) | p = 0.043 | -0.09 |
| 1 | Subgenual /<br>Accumbens | Pre-PBM | Frequency<br>40 > 10 (Hz) | p = 0.034 | +0.12 |
| 2 | Posterior Superior /<br>Temporal Sulcus | Pre-Post | Frequency<br>40 < 10 (Hz) | p = 0.045 | -0.07 |
| 3 | Posterior Cingulate /<br>DMN | Pre-PBM | Frequency<br>40 < 10 (Hz) | p = 0.029 | -0.16 |
| 1 | Subgenual /<br>Accumbens | Pre-PBM | ITA<br>L < I < D | p = 0.015 | -0.05 |
| 3 | Posterior Cingulate /<br>DMN | Pre-Post | ITA<br>L < I < D | p = 0.035 | -0.15 |
| 3 | Posterior Cingulate /<br>DMN | Pre-PBM | Sex<br>Female > Male | p = 0.047 | +0.14 |
| 1 | Subgenual /<br>Accumbens | Pre-PBM | Wavelength * Frequency<br>1064 nm * 40 Hz | p = 0.015 | -0.13 |
| 3 | Posterior Cingulate /<br>DMN | Pre-PBM | Wavelength * Frequency<br>1064 nm * 40 Hz | p = 0.033 | -0.40 |
| 3 | Posterior Cingulate /<br>DMN | Pre-Post | Wavelength * ITA<br>1064 nm * ITA | p = 0.034 | -0.20 |
| 4 | Stimulus Site /<br>Prefrontal Cortex | Pre-Post | Wavelength * ITA<br>1064 nm * ITA | p = 0.046 | +0.21 |

**b) %CBF**
| ROI | Location | Effect | Significance FDR<br>corrected p value | Effect Size |
| --- | --- | --- | --- | --- |
| 1 | Subgenual / Accumbens | Wavelength<br>1064 < 808 (nm) | p = 0.009 | -2.12 |
| 2 | Posterior Superior /<br>Temporal Sulcus | Wavelength<br>1064 < 808 (nm) | p = 0.038 | -0.56 |
| 3 | Posterior Cingulate / DMN | Wavelength<br>1064 < 808 (nm) | p = 0.002 | -1.06 |
| 4 | Stimulus Site / Prefrontal<br>Cortex | Wavelength<br>1064 < 808 (nm) | p = 0.048 | -0.44 |
| 1 | Subgenual / Accumbens | Irradiance<br>100 < 150 > 200 (mW/cm<sup>2</sup>) | p = 0.049 | -0.01 |
| 2 | Posterior Superior /<br>Temporal Sulcus | Irradiance<br>100 < 150 < 200 (mW/cm<sup>2</sup>) | p = 0.019 | 0.86 |
| 2 | Posterior Superior /<br>Temporal Sulcus | ITA<br>L > I > D | p = 0.046 | 0.84 |
| 2 | Posterior Superior /<br>Temporal Sulcus | Sex<br>Male > Female | p = 0.036 | 0.98 |
| 4 | Stimulus Site / Prefrontal<br>Cortex | Sex<br>Male > Female | p = 0.047 | 0.44 |

The differences between Pre-PBM and Pre-Post effects largely reflect the underlying temporal profiles of each ROI’s response to stimulation. ROI 1, which exhibited a block-shaped response that rose during stimulation but returned to baseline post stimulation, showed significant effects only in the Pre-PBM comparisons. This is expected, as the post-stimulation period provided no additional signal beyond baseline. In contrast, ROIs 2-4 displayed a ramp-like trajectory in which the %BOLD response continued to increase even after stimulation. As a result, these regions produced significant findings in both Pre-PBM and Pre-Post t-tests, capturing not only the immediate effects of stimulation but also the lingering cumulative post-stimulation response. Moreover, the pattern of statistical significance across t-tests mirrors the temporal dynamics of each ROI. The anatomical distance from the skull to the cortical surface, calculated per participant, was also included in the LME model. Across all ROIs and conditions, the skull-to-cortex distance did not produce any significant effects.

### Biophysical Modeling

Time-window average %BOLD and %CBF values for all scans (n=180) represented by scatter plots are shown below (**Figure 4a)**. In general, regions showing increased BOLD responses demonstrated proportional increases in CBF. It can be seen that the BOLD-CBF slope decreases with advancing time both during and after the PBM stimulus **(Figure 4b)**.

**Figure 4.**
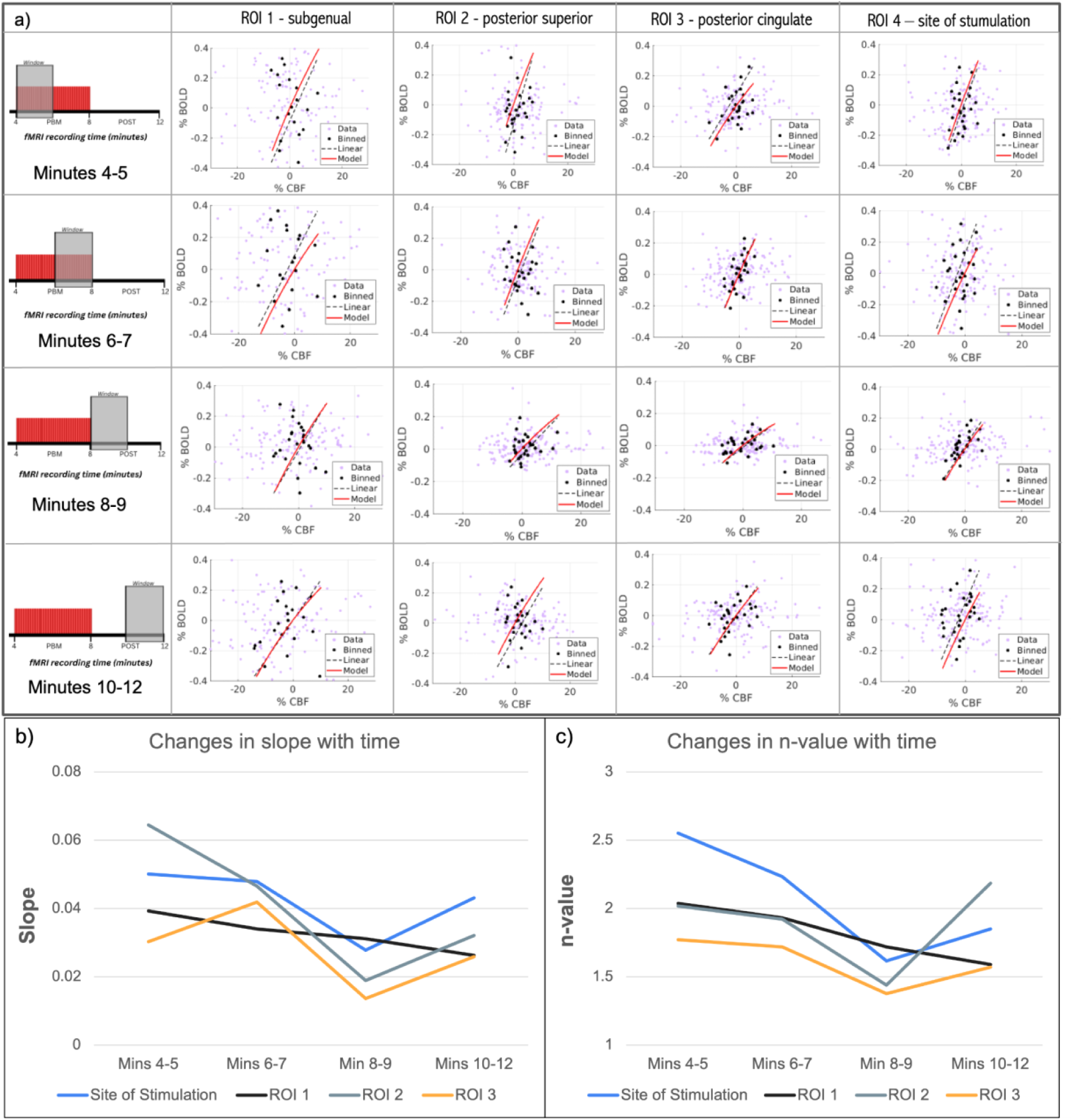
The relationship between BOLD and CBF responses across ROIs was evaluated using a neurovascular coupling model derived from the average BOLD and CBF response. (a) Scatter plot of the temporal mean BOLD and CBF responses from each scan of each subject, derived from the deoxyhemoglobin dilution model (Eq.1). The data were stratified into bins of 6 data points, and each dot represents the mean of one bin. (b) BOLD-CBF slope for each ROI, in time windows. c) n-value variations with time window for all ROIs.

The n-values were estimated by modeling the measured BOLD using the CBF responses, and reflects the relationship between CBF and CMRO_2_. The average n-value was calculated to be 2.04 +/-0.59 (ROI4), 1.78 +/- 0.41 (ROI1), 2.12 +/- 0.76 (ROI2) and 1.61 +/- 0.23 (ROI3). The variations in the estimated n-value **(Figure 4c)** follow those of the BOLD-CBF slopes.

## Discussion

Currently, a major barrier in achieving broader clinical translation of tPBM is the limited understanding of its physiological mechanisms and the associated dose dependence. In this work, we utilized BOLD- and CBF-fMRI to characterize the real-time regional cerebral hemodynamics to tPBM across various stimulation parameters, different sexes and skin tones. Our main findings are: (1) the fMRI response to tPBM is not constrained to the site of stimulation, but is instead distributed across brain regions; (2) the tPBM-evoked fMRI response did not follow a uniform temporal profile; in some regions it closely tracked the stimulation period, whereas in others in remained elevated after stimulation; (3) parameter associations were regionally dependent; with varied responses for BOLD and CBF; (4) the response also depended on skin tone and sex; (5) the BOLD fMRI response to tPBM generally suggests a contribution by CBF, but the BOLD-CBF relationship varied with stimulation time period. These results were found to be independent from any thermal mechanism as demonstrated in our MR thermometry section in Results and Supplementary Materials.

### Spatial distribution of BOLD response to tPBM

Our results show a BOLD response at the site of stimulation, consistent with the current literature, however, this response was modest. Greater BOLD responses are found in other ROIs, including the subgenual (ROI1), the posterior superior temporal sulcus (ROI2, the largest BOLD response) as well as the posterior cingulate (ROI3). We speculated that the increases in the BOLD fMRI signal are mediated through similar mechanisms as the increase in functional connectivity [19], with an enhanced CBF and modulated cerebral oxygenation.

The posterior superior/temporal sulcus regions are highly connected with the site of stimulation [39], forming part of the DMN. These connections may facilitate the sequential activation of distal nodes following prefrontal stimulation, a phenomenon previously also reported in transcranial magnetic stimulation [40]. Furthermore, in our recent work, we dissect the sequence of this propagation [41]. We noted that BOLD increases appeared in the medial and lateral prefrontal cortices immediately upon tPBM onset, with subsequent activation emerging in regions associated with the DMN over the next 40-100s, suggesting that tPBM engages distributed functional networks that evolve over time, rather than acting locally at the site of stimulation.

### Temporal dynamics of BOLD response to tPBM

As the neurovascular coupling status of PBM is unknown, we used a model-free approach. This analysis yielded two distinct temporal patterns that were region-dependent, as shown in **Figure 3**. ROI 1 demonstrates a block pattern corresponding largely to the onset and offset times of our light stimulus. This shape has been noted in previous BOLD-fMRI [22] and functional connectivity studies [19], but is inconsistent with the vast body of fNIRS literature, which report a sustained post-stimulus hemodynamic response [42], [43]. Interestingly, the BOLD responses in ROIs 2-4 all exhibit a more sustained BOLD response shape, that has also been seen through oxyhemoglobin concentration with fNRIS [44]. Moreover, a change in deoxyhemoglobin concentration at 3 Tesla corresponds approximately 0.055% of the baseline BOLD signal change, consistent with the magnitude of %BOLD responses observed in our study [45].

The fNRIS literature suggested that the combination of increased mitochondrial activity and NO-mediated vasodilation leads to elevated blood flow and oxygen delivery, producing a sustained increase in Δ[HbO] that continues post-stimulus. This was indeed the case for our BOLD responses at the site of stimulation.

Unlike in fNIRS, fMRI allows measurements from sites well beyond the site of stimulation, where more distal brain regions follow a different response shape that was previously reported by Zhao et al. [22], albeit using continuous PBM and in the frontal lobe. This response is more rapid to recover to baseline and suggests the participation of a transient activation of ion channels. These channels are often triggered by longer wavelengths (+ 980 nm), [46], [47] [48] reflective of our response in ROI 1 showing significant BOLD activation at 1064 nm compared to 808 nm, as well as the 1064.17 nm laser used by Zhao et al. tPBM can trigger responses of transient receptor potential (TRP) channels, which can increase metabolic activity and release vasoactive messengers which by extension increase blood flow [49]. These channels operate on a millisecond timescale, opening and closing rapidly, but their collective activation during PBM produces sustained neuronal and vascular effects, measurable by fMRI. Further investigations are needed to clarify the sources of these different BOLD temporal dynamics.

### BOLD response dose dependence

In human PBM research, there is currently no response-driven dose optimization. More recently, there have been efforts to model the light penetration into the human brain [24], [25], [26]. These simulation results suggest that the 808nm wavelength provides deeper penetration than the 1064nm and 670nm wavelengths. The simulations also predicted a linear dependence of energy deposition on irradiance, contrasting cell-culture studies which have shown a bi-phasic dependence [50]. Nonetheless, there are major limitations as to how brain models and cell culture findings emulate in vivo responses. The current work shows a clear dose dependence for the tPBM BOLD response, which was also region-dependent. When targeting the temporal sulcus and posterior cingulate, 808nm produced the largest BOLD amplitude, while in the subgenual ROI, 1064 nm produced the highest BOLD temporal responses (**Figure S4a, Table 3a**). Irrespective of the mechanism, it is likely that 808 nm is not always the optimal irradiation wavelength.

Moreover, our results suggest a biphasic irradiance dependence, in ROI 1, the response peaked at the intermediate irradiance of 150 mW/cm², but not in ROI 4. In ROI 3, which is mainly associated with the DMN and is further from the source of stimulation, the response scaled with irradiance (**Figure S4b**).

The spatial variations in dose dependence are also dependent on pulsation frequency. For instance, the subgenual cortex (ROI1), which demonstrates the block pattern temporal dynamics at 40 Hz tPBM, is primarily associated with theta and alpha rhythms [51]. Therefore, higher pulsation frequency may disrupt neuronal connections in this ROI, producing transient changes rather than sustained increases (**Figure S4c**). In contrast, both the temporal sulcus (ROI2), and the posterior cingulate cortex (ROI3), showed a sustained ramp response in the BOLD signal, but more pronounced at 10 Hz. Of course, beyond its dominant baseline rhythm, each region’s BOLD response amplitude and shape are also likely driven by its neurovascular interactions.

Lastly, the effects of wavelength and frequency interacted. We noted that both ROI 1 (subgenual) and ROI 3 (posterior cingulate) showed a reduced %BOLD response at a higher pulsation frequency, but only at 1064 nm. In ROI 1, however, the coupling of higher pulsation frequency and longer wavelength produced a negative effect size, similar to ROI 3. These patterns call attention to the combination of parameters; the effects of which are not necessarily additive.

### Biological factors

Regarding skin tone, the significant effect of ITA on PBM response was observed across multiple ROIs, in principle echoing our simulations [26] and underscore the importance of accounting for individual melanin differences in PBM. As predicted by simulation, in ROI 2 (temporal sulcus), individuals with darker skin exhibited lower CBF responses. However, this was not true in ROI 1 (subgenual) and ROI 3 (posterior cingulate), where individuals with darker skin tones exhibited greater %BOLD responses (**Figure S3d**). It is likely regions proximal and distal to the light source may undergo somewhat different physiological responses. ITA also interacted with various light parameters. For instance, in ROI 4 (stimulus site), the combination of 1064nm and higher ITA increased the BOLD response, however, neither wavelength nor ITA alone showed a significant effect in this region.

The deviations between these findings and Monte Carlo simulations [26] highlight the importance of in vivo experiments in determining tPBM dose dependence. One possible mechanism for darker skin eliciting higher BOLD responses is that the lower photon flux in darker skin elicits a compensatory neurovascular amplification, analogous to the hormetic/biphasic responses widely observed with irradiance [50], yielding a disproportionately large hemodynamic effect once a biological threshold is crossed. Alternatively, individuals with darker skin may simply have different baseline metabolic profiles in these mood- and DMN-related networks. Whatever the cause, the data indicates that darker skin does not uniformly reduce tPBM efficacy and may, in certain networks, enhance it.

A significant sex effect was also observed in the responses, but only in ROI 3, with a higher BOLD response noted in females (**Figure S3e**). This difference could be driven by anatomical factors such as skull thickness, but unlikely given that women typically exhibit greater frontal bone thickness [52]. A more likely source is that women also exhibit higher resting CBF [53] and stronger neurovascular regulation compared to men. Lastly, the anatomical distance from the skull to the cortical surface, which could also differ between sexes and individuals, did not significantly contribute.

### The role of CBF

Despite the common assumption that a greater BOLD response is more desirable, the interpretation of the BOLD response is impossible without an understanding of the underlying physiological signals (e.g. CBF and CMRO_2_) This study is the first to show evidence of a PBM-driven CBF response across multiple brain regions in real time using MRI. In all regions, we see an increase in %BOLD immediately after the onset of stimulation, reflecting an initial hemodynamic response to neural activity, whether this response is sustained past the stimulation period or returns to baseline. The two distinct temporal patterns of BOLD responses suggest two scenarios of neurovascular interactions, as detailed in **Figure 5**.

**Figure 5.**
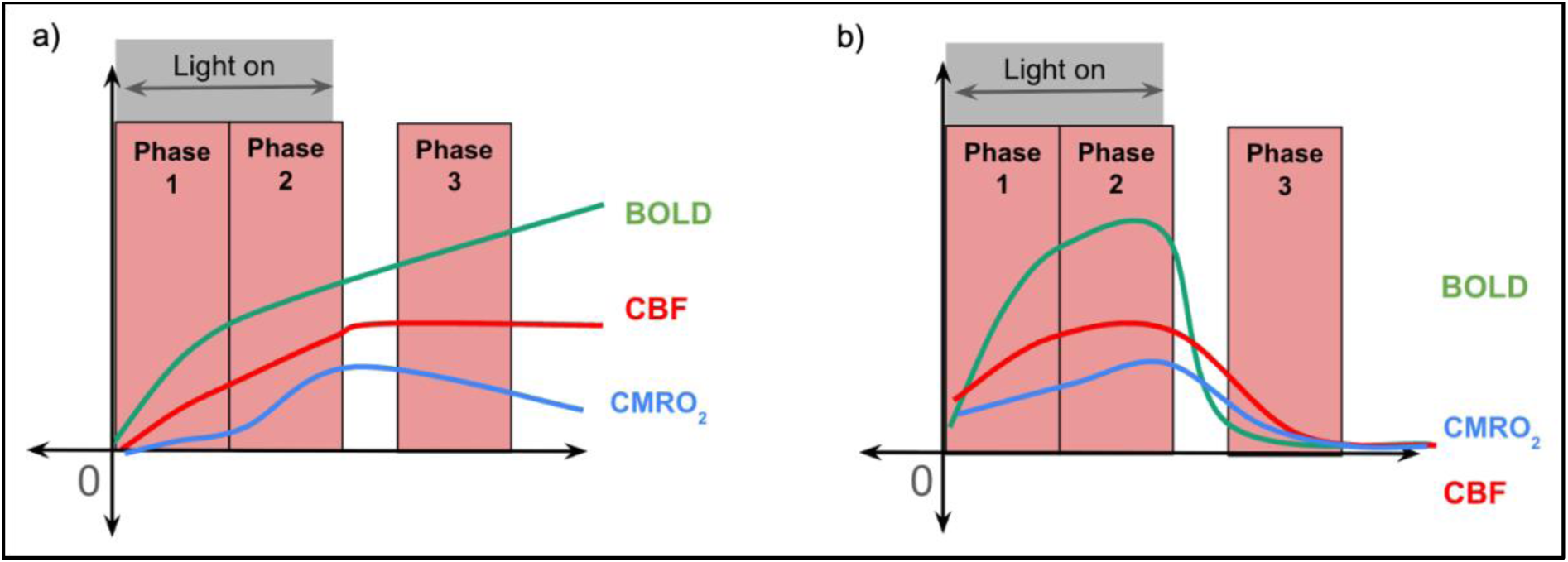
Probable hemodynamic and metabolic responses to tPBM across (a) scenarios 1 and (b) scenario 2. Temporal evolution of the BOLD signal (green), CBF signal (red) and CMRO₂ (blue) shown across the PBM stimulus (grey region - phase 1 and 2) and post stimulus recording (phase 3). The steep BOLD-CBF slope during the first two minutes of stimulation suggests a substantial increase in local blood oxygenation that may arise from a CBF increase. This early phase likely reflects robust arterial dilation and is consistent with the past observations of increased Δ[HbO] using fNIRS [13].

#### Scenario 1: ROIs 2, 3 and site of stimulation (ROI4)

In ROIs 2, 3 and the site of stimulation, the BOLD response persistently climbed, not returning to baseline even after stimulus cessation (**Fig. 5a**). This rise in the BOLD signal would have to be supported by a rise in CBF that is proportionally greater than that of CMRO₂, producing a net rise in oxygenated hemoglobin ΔHbO concentration, and a relative reduction in deoxyhemoglobin (ΔHb) (Phase 1). During the course of the PBM stimulus, there is a gradual n-value and BOLD-CBF slope decrease, suggesting a reduction in the CBF increase relative to the CMRO₂ increase (Phase 2). After the stimulation ends, CMRO₂ gradually returns to baseline (Phase 3), consistent with previous reports of similar behaviour in oxidized CCO concentration [54]. However, CBF remains elevated due to delayed vascular recovery and sustained vasodilation. This is also consistent with previous fNIRS reports of ΔHbO behaviour [13], and is reflected in the rebound in the n-value post stimulation (**Figure 4b, 4c)**.

The change in coupling between CBF and CMRO₂ (a declining n-value during stimulation followed by a rise in n-value post-stimulation) suggests that CBF and CMRO_2_ responses may not be fully coupled. Regionally, these dynamics likely reflect each region’s baseline ATP, oxygen and glucose metabolism state and electrophysiological profile. Regions that consume more oxygen at baseline or exhibit more high-frequency neural activity (such as gamma rhythm), may show stronger or more sustained hemodynamic effects to tPBM. As tPBM boosts mitochondrial activity and CBF, if a region is already metabolically active, the hemodynamic effect is amplified. In this regard, both the posterior superior temporal sulcus (ROI2) and the prefrontal cortex (ROI4), have a high resting metabolic rate [55], and are dominated by beta and gamma oscillations [56]. The posterior cingulate (ROI3) is part of the DMN, demonstrating high resting glucose and oxygen consumption [57,58]. tPBM in these regions may therefore produce this sustained hemodynamic response, as tPBM further enhances metabolism, when the stimulation ends, the metabolic demand returns to baseline, but CBF remains elevated. This sustained CBF may reflect the involvement of an alternative vascular pathway independent of CMRO₂, or serve to remove metabolic waste generated during the period of heightened oxidative metabolism.

#### Scenario 2: ROI 1

The BOLD response in ROI 1 follows a more conventional trajectory that resembles the block design. There is no maintained post-stimulus BOLD response, and no rebound in the n-value (Phase 3). This suggests that the increased CCO oxidation fuels neuronal activity, leading to the appearance of a tighter neurovascular coupling. Nonetheless, like in Scenario 1, both the BOLD-CBF slope and modeled *n*-value steadily decreased over time (Phase 1-3) (**Figure 4b, 4c)**. This could reflect a gradual rebalancing between flow and metabolism as the brain enters a new neurovascular steady state. These findings are consistent with results from Zhao et al. [22].

The subgenual region is associated with high oxidative metabolism at rest and elevated baseline CMRO₂. Its resting electrophysiology is dominated by theta and alpha rhythms. These properties likely limit the region’s ability to sustain prolonged BOLD elevations, while tPBM enhances mitochondrial activity and transiently increases CMRO₂, the combination of high baseline metabolic demand and slower oscillatory dynamics resulting in a block-shaped response that peaks and saturates during stimulation, then declines post stimulation. It is possible that the subgenual’s distinctive structural and functional organization [59], particularly its role as an integrated pathway for regulation [60,61] clears metabolic waste generated during the period of heightened oxidative metabolism, allowing CBF to return to baseline post-stimulus, in contrast to other regions where sustained blood flow persists.

Lastly, baseline differences in aerobic metabolism may underlie these regional differences [62]. The FAM210B protein, which controls lactate production as a marker of anaerobic metabolism [63], exhibits substantial variability across different brain networks. ROI 1 is characterized by the lowest FAM210B concentration of all ROIs [64]. In the context of our results, this low lactate level could underlie the lack of the apparent neurovascular decoupling that’s needed for a sustained post-stimulus BOLD response. In addition, a recent study mapped the mitochondrial respiratory capacity (MRC) across the brain, [65] which reflects the mitochondria’s maximum ability to consume oxygen for oxidative phosphorylation, an indicator of how efficiently a cell can use this oxygen to produce ATP. In post-hoc analysis, we compared the MRC to BOLD dynamics in each of our ROIs. The posterior superior temporal sulcus (ROI2) and the posterior cingulate (ROI3) exhibited the highest average MRC values, 1.053 and 1.039 respectively, coinciding with greater BOLD responses post-stimulus. The average MRC for the subgenual (ROI1) was 0.945, while the site of stimulation (ROI4) had the lowest value of 0.897. This line of evidence further supports the modulation of the PBM BOLD response by baseline metabolic mechanism, which differs across brain regions.

### Network-propagation of BOLD response

PBM enhances mitochondrial activity, which increases the neuronal energy demand. In response, astrocytes can be engaged in lactate production to increase energy production. Astrocytic activity is coupled to the vascular response through calcium signaling [66], which, propagating through the astrocytic gap junctions, could contribute to the network-like propagation of the PBM response [67] [68].

The network aspect of the response is consistent with that of other non-invasive brain stimulation modalities such as TMS and transcranial electrical stimulation. However, while these patterns primarily modulate neuronal excitability, emerging techniques such as PBM and transcranial focused ultrasound show evidence of directly influencing CBF [69]. Thus, there may be overlapping but also distinct mechanisms through which different stimulation approaches engage brain networks.

### Limitations

This study evaluates the immediate effects of a 4-minute tPBM stimulation. While these acute changes in BOLD and CBF provide insight into the brain’s hemodynamic response, it may not reflect long-lasting neuromodulatory changes that could emerge with repeated tPBM stimulations or prolonged light exposure. Data collection was also limited to four minutes of post-stimulation acquisition, preventing characterization of delayed BOLD and CBF responses over a longer period of time. Moreover, CBF measurements have an inherently low signal-to-noise ratio, which may have reduced the sensitivity to small or transient changes in perfusion, and without direct metabolic readouts can constrain the physiological interpretation.

## Conclusion

This study demonstrates that tPBM can modulate both the BOLD and CBF response, across various cortical structures in the human brain through a single-source laser. Stimulation parameters play a key role in dictating the amplitude of the response. Regions including the posterior superior temporal gyrus and the posterior cingulate show that 808nm is more effective at enhancing the BOLD and CBF response compared to structures such as the subgenual which benefits from longer wavelengths, such as 1064nm. Skin tone modulated the magnitude of the BOLD response, with darker skin causing a reduced response. 10Hz frequency modulated the response in the posterior superior temporal gyrus and the posterior cingulate compared to 40Hz. The BOLD response dynamics also varied across regions, suggesting that region-specific neurovascular and metabolic profiles drive the PBM response. To our knowledge, this is the first study to utilize fMRI to map how individual tPBM stimulation parameters modulate BOLD and CBF responses in vivo. We provide the first evidence that distinct stimulation parameters produce dissociable real-time temporal BOLD response profiles, indicating that dose manipulation alters not only the magnitude of the response but also the dynamics. Future work could implement these dosing parameters and provide repeated stimulation to various participant populations.

## Acknowledgments

We wish to thank our funding supporters; the Ontario Centre for Innovation and the Natural Sciences and Engineering Research Council of Canada and we are grateful for financial donations from Ms. Linda Reed. Furthermore, we acknowledge the research insights provided to us by Corrado Calì and Marjorie Dole.

## Author Contribution Statement

Conception and Design - H.V.L and J.J.C; Data Acquisition - H.V.L, X.Z.Z and J.X.C; Data Analysis and Interpretation - H.V.L and J.J.C; Drafting the Article, Critical Revision, Project Administration - J.J.C; Final Approval H.V.L, X.Z.Z, J.X.C and J.J.C; Resource Contribution - J.J.C

## Statements and declarations

### Ethical considerations

This study was approved by the Baycrest Research Ethics Board (REB# 20-28) on October 19th, 2020.

### Consent to participate

All participants provided written informed consent prior to participating.

### Consent for publication

All participants provided written informed consent for publication prior to participating.

### Declaration of conflicting interest

The authors declared no potential conflicts of interest with respect to the research, authorship, and/or publication of this article.

### Funding statement

The authors disclosed receipt of the following financial support for the research, authorship, and/or publication of this article: This work was supported by the Ontario Centre for Innovation and the Natural Sciences and Engineering Research Council of Canada as well as financial donations from Ms. Linda Reed.

### Data availability

The datasets generated during and/or analyzed during the current study are not publicly available as they include participant-level clinical data, but are available from the corresponding author on request.

## Supplementary Materials

### S.1 Methods - Independent Component Analysis (ICA) and Dual Regression

FSL MELODIC is a computational technique designed to distinguish meaningful signal from noise and other variability in neuroimaging data by identifying statistically independent components. The software applies a two-stage decomposition process: first, principal component analysis (PCA) is used to reduce the complexity of the dataset by preserving the most relevant sources of variation; second, ICA is performed to extract spatially and temporally independent components without prior assumptions about the signal sources. By leveraging ICA, FSL MELODIC provides an unbiased approach to decomposing complex neuroimaging data, allowing for the identification of task-related and resting-state networks even in the absence of a predefined model. Thus, this method was used to identify the possible hemodynamic responses of PBM. An overview of the fMRI analysis approach using FSL melodic, and subsequent dual regression is shown in **Figure S.1.**

**Figure S1.**
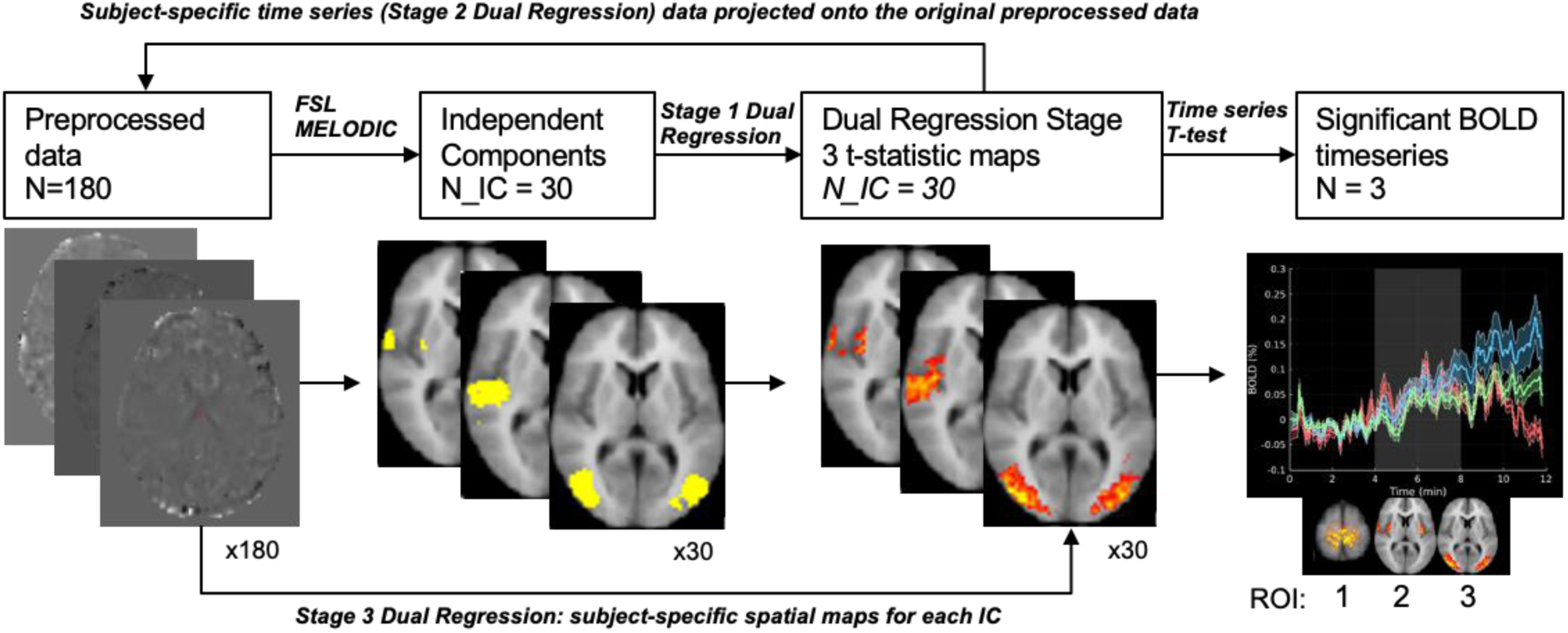
Overview of fMRI analysis approach. Independent component analysis was followed by dual regression for generating the regions of interest (ROIs). All resulting independent components (ICs) were then t-tested for significance in terms of their BOLD signal contrast (both during stimulus and post-stimulus) relative to the pre-stimulus baseline, and the surviving ICs were selected as the final ROIs.

The dual regression model consists of three stages, generating statistical maps and time series for each independent component, allowing for subject-specific characterization of BOLD dynamics and spatial specificity.

i. In the first stage, the group-level ICA spatial maps from MELODIC were regressed onto each participant’s 4D fMRI data using a spatial multiple regression approach. This step extracts subject-specific time series corresponding to each independent component.
ii. In the second stage, these time series are used as regressors in a voxel-wise multiple regression analysis, producing participant-specific spatial maps that reflect the contribution of each ICA component at the individual level.
iii. The third stage involves performing statistical comparisons across subjects or experimental conditions using non-parametric permutation testing, allowing for the identification of significant differences in functional network activity. This approach enables a subject-specific characterization of BOLD dynamics while accounting for inter-individual variability.

After completing the third stage of dual regression, the resulting t-statistic maps were overlaid onto the BOLD-normalized and CBF-normalized time series to extract subject-specific time courses based on the various stimulation parameters. These parameters included three irradiance levels (100, 150, and 200 mW/cm^²^), two wavelengths (808 nm and 1064 nm), two stimulation frequencies (10 Hz and 40 Hz), and three melanin levels representing light, intermediate, and dark skin tones. This approach allowed us to systematically assess how each experimental condition influenced the BOLD and CBF signal fluctuations over time.

After completing the third stage of dual regression, the resulting t-statistic maps were overlaid onto the BOLD-normalized and CBF-normalized time series to extract subject-specific time courses based on the various stimulation parameters. These parameters included three irradiance levels (100, 150, and 200 mW/cm^²^), two wavelengths (808 nm and 1064 nm), two stimulation frequencies (10 Hz and 40 Hz), and three melanin levels representing light, intermediate, and dark skin tones. This approach allowed us to systematically assess how each experimental condition influenced the BOLD and CBF signal fluctuations over time.

### S.1.2 Methods - the individual typology angle (ITA)

The ITA is calculated as (Eq. 1),

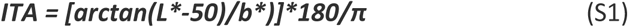

where L represents luminance ranging from black (0) to white (100), and b* ranging from yellow to blue (12). The higher the ITA, the lighter the skin. The ITA skin color types are typically classified into six ranges: very light (>55°), light (41° to <55°), intermediate (28° to <41°), tan (10° to <28°), brown (-30° to <10°) and dark (<-30°). Zero and wWhite calibration were performed before each use to ensure accuracy of the calculations. Six 8mm diameter measurement samples were taken of the right forehead of each subject prior to tPBM laser placement and the average was computed into the individual melanin groupings.

### S.1.3 Methods - MR Thermometry

We utilized an MR thermometry sequence that exploits the temperature sensitivity of the resonance frequency. The MR thermometry sequence followed the same stimulus paradigm as the BOLD-CBF-fMRI sequence shown in **Figure 4**. The temperature was measured in the illuminated region shown in **Figure S1**. The illuminated region was delineated following guidance from our previous work using Monte Carlo simulations to map light propagation [26]. The MRI phase and magnitude images were processed to estimate temperature changes induced by tPBM using the Proton Resonance Frequency (PRF) shift method. A custom preprocessing script was used for, brain extraction, motion correction, drift correction, artifact removal (4 standard deviations above the average), phase radian conversion, unwrapping corrected for phase aliasing to produce a continuous phase map. Temperature changes (ΔT) were then calculated as follows, with all symbols defined in **Table S1**.:

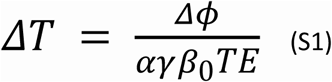

Temperature changes were calculated on a frame by frame basis, using Equation 5 and the following parameters shown in **Table 4**.

**Table S1:** MR thermometry parameters.

| Symbol |  | Value | Units |
| --- | --- | --- | --- |
| $\Delta T$ | Frame by frame temperature change | | $^{\circ}\text{C}$ |
| $\Delta$ | Phase change from baseline | 0 : 4095 | radians |
| $\gamma$ | Gyromagnetic ratio of hydrogen | $2\pi \cdot 42.58 \times 10^6$ | rad/s/T |
| $\alpha$ | PRF thermal coefficient | $0.01 \times 10^{-6}$ | ppm/$^{\circ}\text{C}$ |
| $B_0$ | Main magnetic field strength | 3 | Telsa |
| TE | Echo time | 0.00738 s | seconds |

Temperature time courses were analyzed separately for the two frequency groups; 10Hz (N=16) and 40Hz (N=14). The temperature of each subject was recorded per frame, and then the mean temperature change was calculated for group-level response. The standard error was also calculated to provide an estimate of variability between subjects. A paired sample t-test was performed to compare the mean temperature during the pre-stimulus recording (minutes 0-4) and PBM stimulus (minutes 4-8), to determine if the mean temperature during PBM was significantly different than the mean temperature during the pre-stimulus recording period.

**Figure S2:**
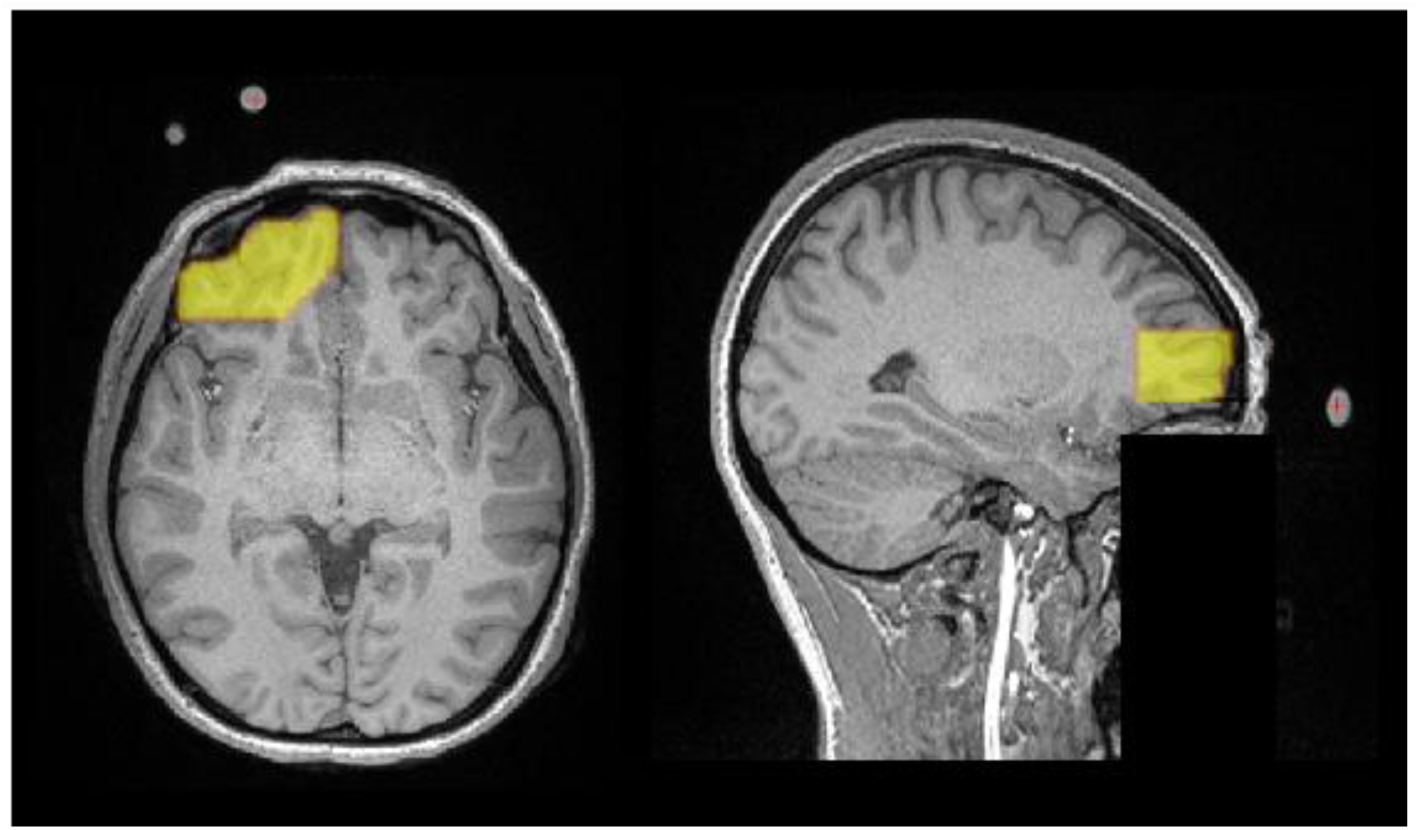
The vitamin E capsules were used as fiducial markers to indicate the laser location. Neural tissue in closest proximity to these markers was segmented, for each participant, and this region was used for MR thermometry analysis.

**Figure S3:**
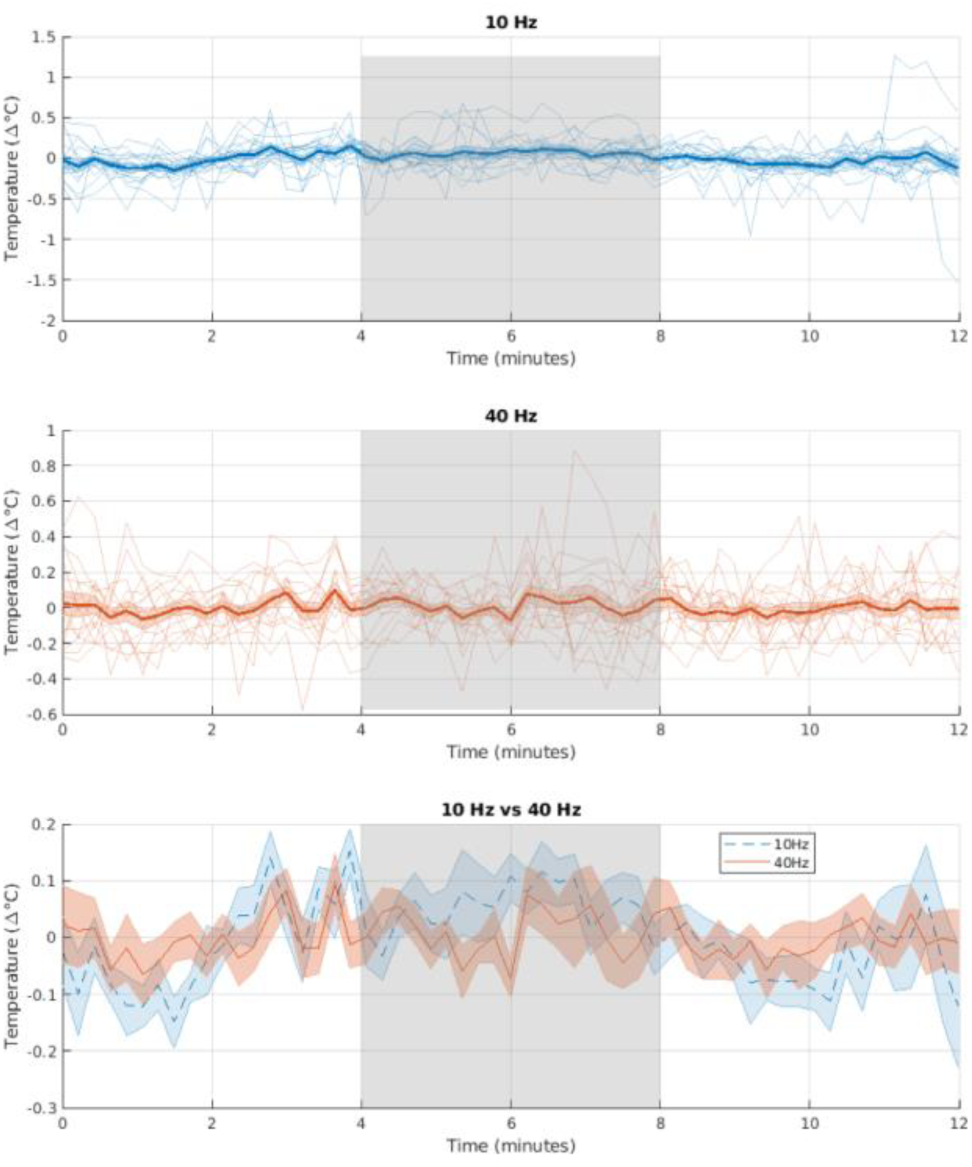
Temperature changes (Δ°C) over time during tPBM at 10 Hz (top), 40 Hz (middle), and their comparison (bottom). Shaded regions indicate the stimulation window (4–8 minutes). Thin lines represent individual subject responses; bold lines show group averages with standard error shading. The comparison plot highlights mean temperature trends for both frequencies, revealing no significant differences in temperature during the stimulation periods.

**Table S2.** Group-level statistical comparison of MR-thermometry-based temperature changes between PBM and the PRE-stimulus recording and standard deviation (STD) for both stimulation frequencies (10 Hz and 40 <u>Hz).</u>.

| Frequency | PRE ($^{\circ}\text{C}$) | PRE STD ($^{\circ}\text{C}$) | PBM ($^{\circ}\text{C}$) | PBM STD ($^{\circ}\text{C}$) | p-value |
| --- | --- | --- | --- | --- | --- |
| 10 Hz | -0.02 | 0.18 | 0.05 | 0.17 | 0.13 |
| 40 Hz | -0.01 | 0.16 | 0.01 | 0.15 | 0.76 |

### S.2.1 Results: Amplitude of the BOLD response by ROI

**Table S3.** Summary of regional %BOLD measurements.

| ROI | Brain regions | Response shape | Mean % BOLD increase during stimulus | Mean % BOLD increase post stimulus | Efficiency (%BOLD /mW.cm<sup>-2</sup>) |
| --- | --- | --- | --- | --- | --- |
| 1 | subgenual / accumbens | Block response | 0.053%<br>SE = 0.04 | 0.023%<br>SE = 0.04 | $2.8 \times 10^{-4}$<br>SE = $1.3 \times 10^{-4}$ |
| 2 | posterior superior / temporal sulcus | Ramp response | 0.021%<br>SE = 0.09 | 0.061%<br>SE = 0.09 | $2.95 \times 10^{-4}$<br>SE = $1.8 \times 10^{-4}$ |
| 3 | posterior cingulate / DMN | Ramp response | 0.002%<br>SE = 0.04 | 0.046%<br>SE = 0.04 | $1.57 \times 10^{-4}$<br>SE = $1.8 \times 10^{-4}$ |
| 4 | stimulus site / prefrontal cortex | Ramp response | 0.004%<br>SE = 0.03 | 0.052%<br>SE = 0.03 | $2.17 \times 10^{-4}$<br>SE = $8.0 \times 10^{-5}$ |

### S.2.2 Results: Linear Mixed Effects Modeling Dose Dependance

**Figure S4.**
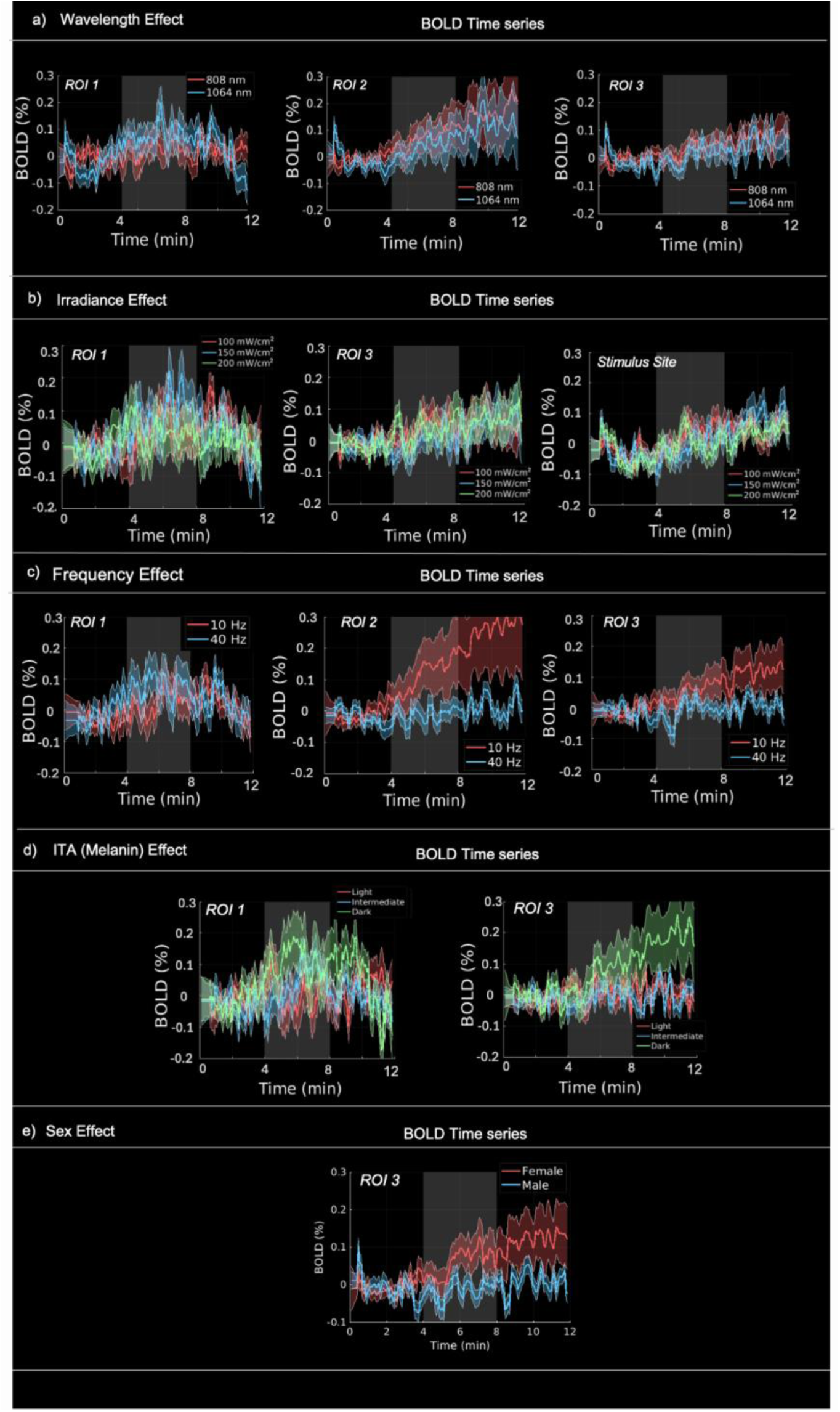
BOLD temporal response magnitudes across stimulation parameters. For each significant parameter association **(Table 4)** within each ROI, the corresponding BOLD time series data was extracted to visualize the differences between values. The tPBM stimulus “on” period (minutes 4:8) is denoted by the shaded grey region. Coloured shaded regions represent the standard error across all datasets.

### S.2.3 Results: Minute by Minute CBF and BOLD responses

The BOLD data derived from the 4 ROIs are averaged across *N* = 180 datasets (4 scans/participant x 45 participants), as shown in **Figure 5**. CBF signal data is inherently noisier compared to BOLD-fMRI signals. We averaged the CBF signal across 1-minute time windows in order to visualize the CBF time course and reduce noise while preserving the actual temporal response. For comparison, we applied the same approach to the BOLD response, as shown in **Figure S3**.

**Figure S5:**
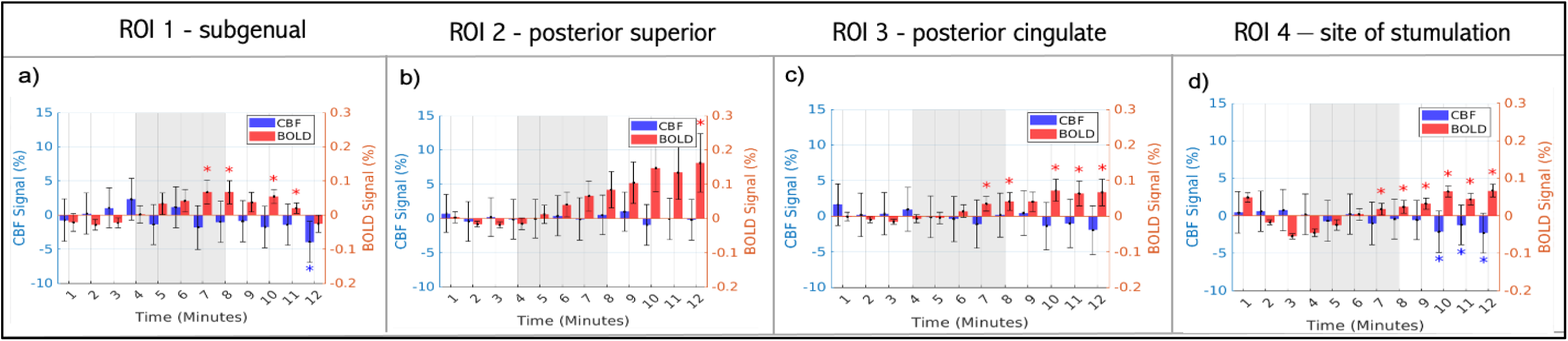
Extracted BOLD (red) and CBF (blue) time course averages for each ROI (a-c) and d) site of stimulation. Each point represents the mean signal for a 1-minute time window across all subjects and scans (n = 180) and for each ROI shown in **Figure 3**. Error bars represent the standard error across all datasets. The PBM stimulus (minutes 4:8) is denoted by the shaded grey region. Asterisks * indicate minutes where the response was significantly different from baseline pre-stimulus (minutes 0:4) based on a two-tailed one-sample t-test across subjects (p < 0.05)

### S.2.4 Results: Linear Mixed Effects Modeling

**Table S4:** BOLD response dose dependence, results from the LME.

| Pre- PBM | ROI 1 - Subgenual / Accumbens |  |  |  |  |  |
| --- | --- | --- | --- | --- | --- | --- |
|  | Variable | Effect Size | DoF | fdr_pvalue | lower bound | upper bound |
|  | Melanin (ITA) | -0.009 | 175 | **0.015** | -0.041 | 0.007 |
|  | Irradiance_150 | 0.015 | 175 | **0.045** | 0.003 | 0.045 |
|  | Irradiance_200 | 0.011 | 175 | **0.045** | -0.028 | 0.030 |
|  | Frequency_40 | 0.022 | 175 | **0.034** | 0.007 | 0.043 |
|  | Wavelength_1064 | 0.025 | 175 | **0.006** | 0.007 | 0.027 |
|  | Sex_Male | -0.013 | 175 | 0.259 | -0.042 | 0.017 |
|  | Distance (Skull to Cranium) | -0.0001 | 175 | 0.915 | -0.002 | 0.002 |
|  | Wavelength_1064: Frequency_40 | -0.031 | 175 | **0.015** | -0.055 | -0.009 |
|  | Wavelength_1064:Melanin (ITA) | -0.009 | 175 | 0.596 | -0.045 | 0.026 |
| Pre- PBM | ROI 2 - Posterior Superior / Temporal Sulcus |  |  |  |  |  |
|  | Variable | Effect Size | DoF | fdr_pvalue | lower bound | upper bound |
|  | Melanin (ITA) | -0.005 | 175 | 0.206 | -0.014 | 0.003 |
|  | Irradiance_150 | -0.005 | 175 | 0.455 | -0.017 | 0.008 |
|  | Irradiance_200 | -0.004 | 175 | 0.455 | -0.017 | 0.008 |
|  | Frequency_40 | -0.011 | 175 | **0.037** | -0.039 | 0.021 |
|  | Wavelength_1064 | 0.005 | 175 | 0.463 | -0.009 | 0.020 |
|  | Sex_Male | 0.004 | 175 | 0.627 | -0.011 | 0.018 |
|  | Distance (Skull to Cranium) | 0.001 | 175 | 0.135 | -0.0003 | 0.002 |
|  | Wavelength_1064: Frequency_40 | -0.015 | 175 | 0.227 | -0.036 | 0.006 |
|  | Wavelength_1064:Melanin (ITA) | 0.012 | 175 | 0.208 | -0.004 | 0.027 |
| Pre- PBM | ROI 3 - Posterior Cingulate / DMN |  |  |  |  |  |
|  | Variable | Effect Size | DoF | fdr_pvalue | lower bound | upper bound |
|  | Melanin (ITA) | 0.003 | 176 | 0.462 | -0.005 | 0.012 |
|  | Irradiance_150 | 0.022 | 176 | **0.041** | -0.030 | 0.002 |
|  | Irradiance_200 | 0.003 | 176 | **0.041** | -0.017 | 0.011 |
|  | Frequency_40 | -0.017 | 176 | **0.029** | -0.018 | 0.006 |
|  | Wavelength_1064 | 0.015 | 176 | 0.101 | -0.001 | 0.030 |
|  | Sex_Male | 0.015 | 176 | **0.047** | -0.004 | 0.029 |
|  | Distance (Skull to Cranium) | 0.0002 | 176 | 0.779 | -0.001 | 0.001 |
|  | Wavelength_1064: Frequency_40 | -0.001 | 176 | **0.033** | -0.047 | -0.004 |
|  | Wavelength_1064:Melanin (ITA) | 0.004 | 176 | 0.433 | -0.016 | 0.024 |
| Pre- PBM | ROI 4 - Stimulus Site / Prefrontal Cortex |  |  |  |  |  |
|  | Variable | Effect Size | DoF | fdr_pvalue | lower bound | upper bound |
|  | Melanin (ITA) | 0.008 | 177 | 0.479 | -0.003 | 0.019 |
|  | Irradiance_150 | -0.009 | 177 | 0.248 | -0.022 | 0.004 |
|  | Irradiance_200 | 0.007 | 177 | 0.248 | -0.005 | 0.020 |
|  | Frequency_40 | -0.007 | 177 | 0.649 | -0.025 | 0.010 |
|  | Wavelength_1064 | 0.007 | 177 | 0.212 | -0.004 | 0.017 |
|  | Sex_Male | -0.004 | 177 | 0.413 | -0.015 | 0.006 |
|  | Distance (Skull to Cranium) | -0.0007 | 177 | 0.239 | -0.002 | 0.001 |
|  | Wavelength_1064: Frequency_40 | 0.009 | 177 | 0.233 | -0.024 | 0.006 |
|  | Wavelength_1064:Melanin (ITA) | 0.000 | 177 | 0.587 | -0.021 | 0.010 |
| Pre-Post | ROI 1 - Subgenual / Accumbens |  |  |  |  |  |
|  | Variable | Effect Size | DoF | fdr_pvalue | lower bound | upper bound |
|  | Melanin (ITA) | 0.007 | 177 | 0.347 | -0.008 | 0.023 |
|  | Irradiance_150 | -0.017 | 177 | 0.183 | -0.043 | 0.008 |
|  | Irradiance_200 | -0.020 | 177 | 0.183 | -0.046 | 0.005 |
|  | Frequency_40 | -0.013 | 177 | 0.473 | -0.043 | 0.017 |
|  | Wavelength_1064 | 0.013 | 177 | 0.219 | -0.008 | 0.034 |
|  | Sex_Male | 0.032 | 177 | 0.201 | -0.004 | 0.067 |
|  | Distance (Skull to Cranium) | 0.0008 | 175 | 0.518 | -0.002 | 0.003 |
|  | Wavelength_1064: Frequency_40 | 0.032 | 177 | 0.411 | -0.010 | 0.073 |
|  | Wavelength_1064:Melanin (ITA) | -0.056 | 177 | 0.387 | -0.009 | 0.025 |
| Pre-Post | ROI 2 - Posterior Superior / Temporal Sulcus |  |  |  |  |  |
|  | Variable | Effect Size | DoF | fdr_pvalue | lower bound | upper bound |
|  | Melanin (ITA) | -0.006 | 175 | 0.322 | -0.017 | 0.006 |
|  | Irradiance_150 | 0.003 | 175 | 0.762 | -0.014 | 0.019 |
|  | Irradiance_200 | 0.001 | 175 | 0.762 | -0.016 | 0.018 |
|  | Frequency_40 | -0.011 | 175 | **0.045** | -0.006 | 0.021 |
|  | Wavelength_1064 | -0.011 | 175 | **0.041** | -0.012 | 0.036 |
|  | Sex_Male | 0.007 | 175 | 0.293 | -0.006 | 0.021 |
|  | Distance (Skull to Cranium) | 0.0009 | 175 | 0.345 | -0.001 | 0.003 |
|  | Wavelength_1064: Frequency_40 | -0.011 | 175 | 0.082 | -0.035 | 0.002 |
|  | Wavelength_1064:Melanin (ITA) | -0.017 | 175 | 0.074 | -0.035 | 0.002 |
| Pre-Post | ROI 3 - Posterior Cingulate / DMN |  |  |  |  |  |
|  | Variable | Effect Size | DoF | fdr_pvalue | lower bound | upper bound |
|  | Melanin (ITA) | -0.016 | 176 | **0.035** | -0.020 | 0.005 |
|  | Irradiance_150 | 0.006 | 176 | 0.282 | -0.005 | 0.019 |
|  | Irradiance_200 | 0.006 | 176 | 0.282 | -0.006 | 0.018 |
|  | Frequency_40 | 0.008 | 176 | 0.255 | -0.006 | 0.022 |
|  | Wavelength_1064 | -0.027 | 176 | **0.033** | -0.029 | -0.006 |
|  | Sex_Male | 0.008 | 176 | 0.216 | -0.005 | 0.023 |
|  | Distance (Skull to Cranium) | 4.99e-05 | 176 | 0.944 | -0.001 | 0.001 |
|  | Wavelength_1064: Frequency_40 | -0.008 | 176 | 0.364 | -0.025 | 0.009 |
|  | Wavelength_1064:Melanin (ITA) | -0.020 | 176 | **0.034** | -0.023 | 0.004 |
| Pre-Post | ROI 4 - Stimulus Site / Prefrontal Cortex |  |  |  |  |  |
|  | Variable | Effect Size | DoF | fdr_pvalue | lower bound | upper bound |
|  | Melanin (ITA) | -0.004 | 177 | 0.440 | -0.013 | 0.006 |
|  | Irradiance_150 | 0.010 | 177 | **0.043** | -0.003 | 0.022 |
|  | Irradiance_200 | 0.011 | 177 | **0.043** | -0.002 | 0.024 |
|  | Frequency_40 | -0.001 | 177 | 0.498 | -0.016 | 0.014 |
|  | Wavelength_1064 | 0.011 | 177 | 0.052 | 0.000 | 0.021 |
|  | Sex_Male | 0.005 | 177 | 0.378 | -0.007 | 0.018 |
|  | Distance (Skull to Cranium) | 0.0003 | 177 | 0.649 | -0.001 | 0.002 |
|  | Wavelength_1064: Frequency_40 | 0.000 | 177 | 0.468 | -0.021 | 0.021 |
|  | Wavelength_1064:Melanin (ITA) | 0.021 | 177 | **0.046** | -0.005 | 0.027 |

**Table S5:**
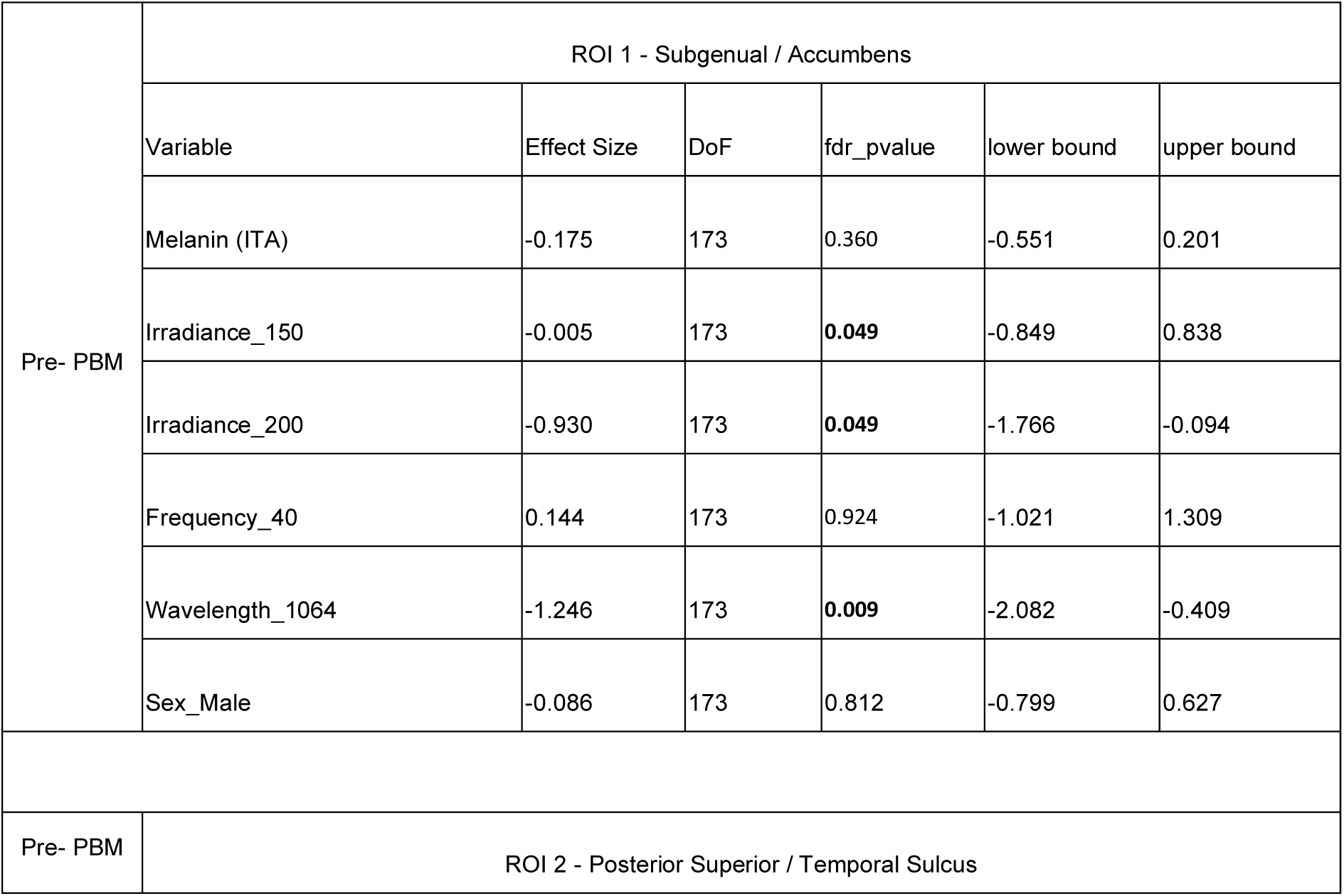
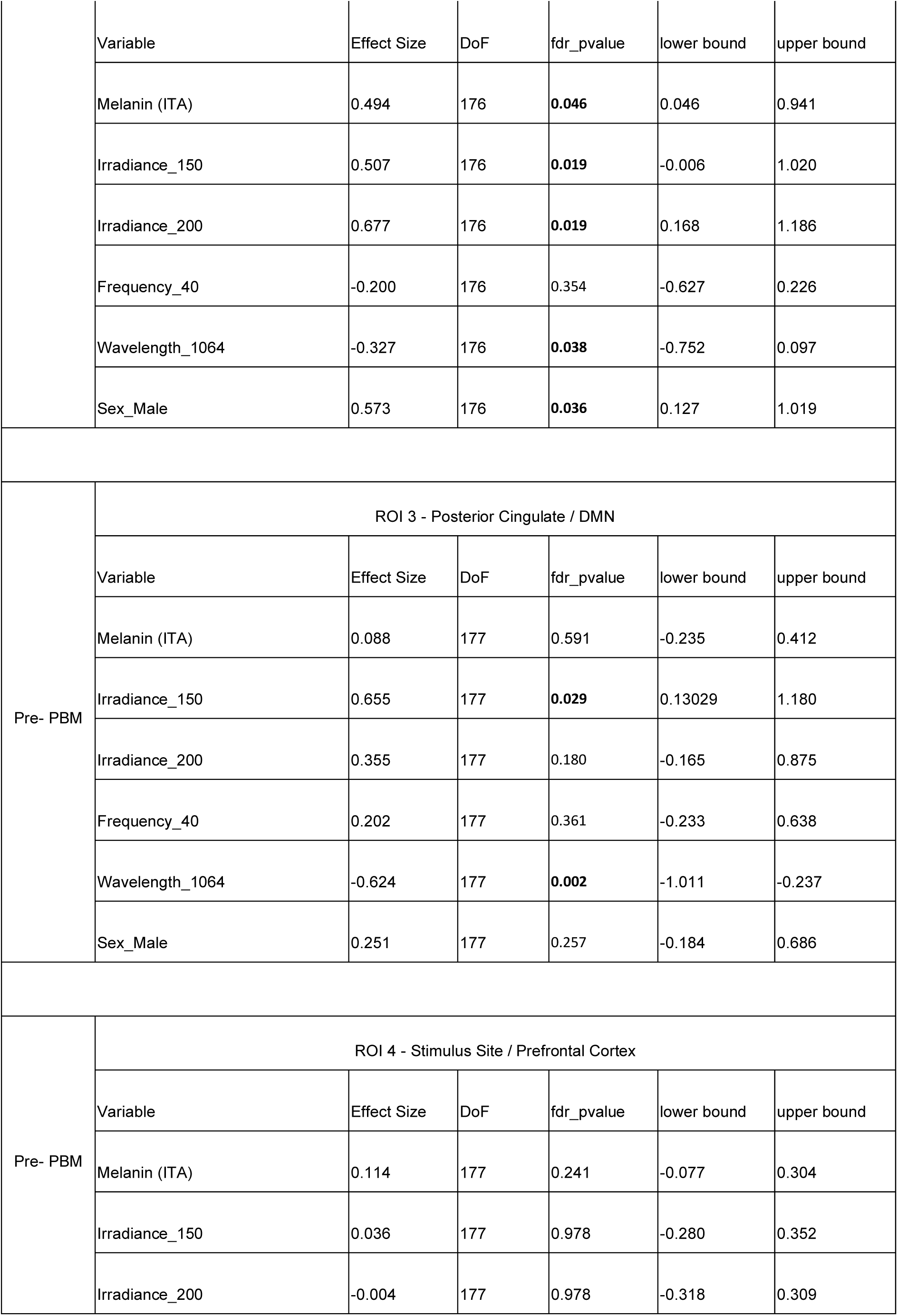
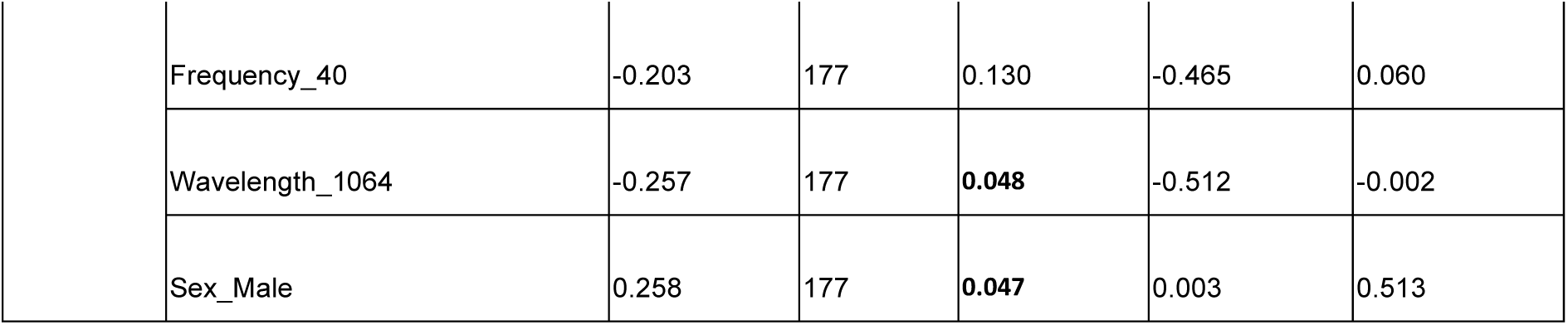
CBF response dose dependence, results from the LME.

|  |  |  |  |  |  |  |
| --- | --- | --- | --- | --- | --- | --- |
| Pre- PBM | ROI 1 - Subgenual / Accumbens |  |  |  |  |  |
|  | Variable | Effect Size | DoF | fdr_pvalue | lower bound | upper bound |
|  | Melanin (ITA) | -0.175 | 173 | 0.360 | -0.551 | 0.201 |
|  | Irradiance_150 | -0.005 | 173 | **0.049** | -0.849 | 0.838 |
|  | Irradiance_200 | -0.930 | 173 | **0.049** | -1.766 | -0.094 |
|  | Frequency_40 | 0.144 | 173 | 0.924 | -1.021 | 1.309 |
|  | Wavelength_1064 | -1.246 | 173 | **0.009** | -2.082 | -0.409 |
|  | Sex_Male | -0.086 | 173 | 0.812 | -0.799 | 0.627 |
| Pre- PBM | ROI 2 - Posterior Superior / Temporal Sulcus |  |  |  |  |  |
|  | Variable | Effect Size | DoF | fdr_pvalue | lower bound | upper bound |
|  | Melanin (ITA) | 0.494 | 176 | **0.046** | 0.046 | 0.941 |
|  | Irradiance_150 | 0.507 | 176 | **0.019** | -0.006 | 1.020 |
|  | Irradiance_200 | 0.677 | 176 | **0.019** | 0.168 | 1.186 |
|  | Frequency_40 | -0.200 | 176 | 0.354 | -0.627 | 0.226 |
|  | Wavelength_1064 | -0.327 | 176 | **0.038** | -0.752 | 0.097 |
|  | Sex_Male | 0.573 | 176 | **0.036** | 0.127 | 1.019 |
| Pre- PBM | ROI 3 - Posterior Cingulate / DMN |  |  |  |  |  |
|  | Variable | Effect Size | DoF | fdr_pvalue | lower bound | upper bound |
|  | Melanin (ITA) | 0.088 | 177 | 0.591 | -0.235 | 0.412 |
|  | Irradiance_150 | 0.655 | 177 | **0.029** | 0.13029 | 1.180 |
|  | Irradiance_200 | 0.355 | 177 | 0.180 | -0.165 | 0.875 |
|  | Frequency_40 | 0.202 | 177 | 0.361 | -0.233 | 0.638 |
|  | Wavelength_1064 | -0.624 | 177 | **0.002** | -1.011 | -0.237 |
|  | Sex_Male | 0.251 | 177 | 0.257 | -0.184 | 0.686 |
| Pre- PBM | ROI 4 - Stimulus Site / Prefrontal Cortex |  |  |  |  |  |
|  | Variable | Effect Size | DoF | fdr_pvalue | lower bound | upper bound |
|  | Melanin (ITA) | 0.114 | 177 | 0.241 | -0.077 | 0.304 |
|  | Irradiance_150 | 0.036 | 177 | 0.978 | -0.280 | 0.352 |
|  | Irradiance_200 | -0.004 | 177 | 0.978 | -0.318 | 0.309 |

|  |  |  |  |  |  |
| --- | --- | --- | --- | --- | --- |
| Frequency_40 | -0.203 | 177 | 0.130 | -0.465 | 0.060 |
| Wavelength_1064 | -0.257 | 177 | **0.048** | -0.512 | -0.002 |
| Sex_Male | 0.258 | 177 | **0.047** | 0.003 | 0.513 |

## Notes

### Competing Interest Statement

The authors have declared no competing interest.

### Summary of Updates

Title Revision and additional work added to the discussion

